# Learning when effort matters: Neural dynamics underlying updating and adaptation to changes in performance efficacy

**DOI:** 10.1101/2020.10.09.333310

**Authors:** Ivan Grahek, Romy Frömer, Mahalia Prater Fahey, Amitai Shenhav

## Abstract

To determine how much cognitive control to invest in a task, people need to consider whether exerting control *matters* for obtaining rewards. In particular, they need to account for the efficacy of their performance – the degree to which rewards are determined by performance or by independent factors. Yet it remains unclear how people learn about their *performance efficacy* in an environment. Here we combined computational modeling with measures of task performance and EEG, to provide a mechanistic account of how people (a) learn and update efficacy expectations in a changing environment, and (b) proactively adjust control allocation based on current efficacy expectations. Across two studies subjects performed an incentivized cognitive control task while their performance efficacy (the likelihood that rewards are performance-contingent or random) varied over time. We show that people update their efficacy beliefs based on prediction errors – leveraging similar neural and computational substrates as those that underpin reward learning – and adjust how much control they allocate according to these beliefs. Using computational modeling, we show that these control adjustments reflect changes in information processing, rather than the speed-accuracy tradeoff. These findings demonstrate the neurocomputational mechanism through which people learn how worthwhile their cognitive control is.

---

Cognitive control is critical for achieving most goals, but it is effortful (Botvinick and Cohen 2014; Shenhav et al. 2017). To decide how to invest control into a task (e.g., writing an essay for a competition), a person must weigh these effort costs against the potential benefits of a given type and amount of control (Manohar et al. 2015; Verguts et al. 2015; Kool and Botvinick 2018). One aspect of these benefits is the significance of the expected outcomes, both positive (e.g., a monetary prize, social acclaim) and negative (e.g., missed revenue, social derision) (Atkinson 1966; Leng et al., 2021). An equally important aspect of the expected benefits of control is the extent to which control *matters* for achieving good outcomes and avoiding bad ones (Frömer et al., 2021a; Shenhav et al., 2021). This can in turn be decomposed into the extent to which higher levels of control translate into better performance (e.g., whether writing a good essay will require substantial or only minimal control resources; *control efficacy*) and the extent to which better performance translates into better outcomes (e.g., whether prizes are determined by the strength of an essay or by arbitrary or even biased metrics unrelated to essay-writing performance; *performance efficacy*). Whereas studies have increasingly characterized the ways in which control allocation is influenced by expected outcomes (e.g., Parro et al., 2018; Leng et al., 2021) and the expected efficacy of control (e.g., as a function of task difficulty; Krebs et al., 2012; Vassena et al., 2014; Chiu and Egner 2019), much less is known about how people estimate and adjust to the perceived efficacy of their performance in a given environment.

We recently showed that when participants are explicitly instructed about how efficacious their performance will be on an upcoming trial, they exhibit behavioral and neural responses consistent with increased control (Frömer et al., 2021a). We had participants perform a standard cognitive control task (Stroop) for potential monetary rewards, and we varied whether obtaining those rewards was contingent on performing well on the task (high performance efficacy) or whether those rewards were determined at random (low efficacy). We showed that people allocate more control when they expect to have high compared to low efficacy, reflected in higher amplitudes of an EEG index of proactive control (the contingent negative variation [CNV]) and in improved behavioral performance. These results demonstrate that participants leverage expectations about the extent to which their performance matters when deciding how much cognitive effort to invest in a task. However, in this work, participants were explicitly cued with the level of performance efficacy to expect on a given trial, and those predictive cues retained the same meaning across the session. Thus, how it is that people learn these efficacy expectations in environments where contingencies are not instructed, and how they dynamically update their expectations as contingencies change, remains unanswered.

Outside of the domain of cognitive control, a relevant line of work has examined how people learn about the factors that determine future outcomes when selecting between potential courses of action. In particular, work in this area has shown that people are able to learn about and update their expectations of the likelihood that a given action will generate a given outcome (action-outcome contingency; Dickinson and Balleine 1995; Moscarello and Hartley 2017; Ly et al. 2019). People preferentially, and more vigorously, select actions that reliably lead to desired outcomes (i.e., the more contingent those outcomes are on the action in question; Liljeholm et al. 2011; Manohar et al. 2017), and work in both animals (Balleine & O’Doherty, 2010) and humans (Norton & Liljeholm, 2020; Dorfman et al., 2021; Ligneul et al., 2022; Morris et al., 2022) has helped to characterize the neural systems that support this process of learning and action selection. However, given the focus on discrete actions and their immediate relationship with outcomes, research into these action-outcome contingencies is unable to capture key aspects that are unique to selection of control states. Most notably, cognitive control signals (e.g., attention to one or more features of the environment) are multidimensional, not immediately observable by either the participant or experimenter, and their relationship with potential outcomes is intermediated by the many-to-many relationship between control states and task performance (Ritz et al., 2022). While there has been research into how people learn to adjust cognitive control signals based on changes in their task environment, here again work has focused on how people adapt to changes in outcomes (e.g., Otto and Daw 2019; Bustamante et al. 2021) and changes in the relationship between control and performance (with increasing task difficulty; Bugg et al. 2011; Nigbur et al. 2015; Bejjani et al. 2018; Jiang et al. 2020). The mechanisms by which people learn about the relationship between performance and outcomes (performance efficacy), and how they adjust their control allocation accordingly, remain largely unexplored.

Here, we seek to fill this gap by studying the mechanisms through which expectations of performance efficacy are formed, updated, and used to guide control allocation. To do so, we extend our previous approach (Frömer et al., 2021a) – which studied how behavioral and neural correlates of control allocation vary when performance efficacy is explicitly cued – to examine how participants learn and adapt their control under conditions where efficacy was un-cued (having to instead be learned from feedback) and gradually varied across a wide range of potential efficacy values over the course of the session. We use computational reinforcement learning models to show that expected efficacy can be learned from feedback through iterative updating based on weighted prediction errors (Sutton and Barto 2018), and model-based single-trial EEG analyses to show that these efficacy prediction errors modulate a canonical neural marker of reward-based learning and behavioral adjustment (Fischer and Ullsperger 2013). We further provide evidence that efficacy estimates learned in this way are used to guide the allocation of control. In our EEG study and a second behavioral study, participants tended to perform better when efficacy was higher. We also provide evidence that a neural marker of control allocation (Schevernels et al. 2014a) tends to increase with increasing model-based efficacy estimates. Using a drift diffusion model (Ratcliff & McKoon, 2008; Wiecki, Sofer, & Frank, 2013), in Study 2 we further show that the performance improvements related to increased performance-efficacy reflect facilitation of task-related information processing (reflected in increased drift rates), rather than changes in the speed-accuracy tradeoff (i.e., thresholds). Taken together, these results show that efficacy estimates can be learned and updated based on feedback, leveraging general cognitive and neural mechanisms of predictive inference.

## Materials and Methods

### Study 1

#### Participants

We recruited forty-one participants with normal or corrected-to-normal vision from the Brown University subject pool. One participant was excluded due to technical issues. The final data set included 40 participants (24 females, 16 males; median age = 19). Participants gave informed consent and were compensated with course credits or a fixed payoff of $20. In addition, they received up to $5 bonus that depended on their task performance ($3.25 on average). The research protocol was approved by Brown University’s Institutional Review Board.

#### Experimental design

In the main task, taking approximately 45 minutes, participants performed 288 Stroop trials (Figure 1A). Each trial started with the presentation of a cue (grey circle) that remained on the screen throughout the trial. After a period of 1500 ms, a Stroop stimulus was superimposed until a response was made or 1000 ms elapsed, at which time it was sequentially replaced with two types of feedback presented for 1000 ms each. Each trial onset was preceded by a fixation cross (randomly jittered between 1000 and 1500ms). Participants responded to the ink color of the Stroop stimulus (red, green, blue, or yellow) by pressing one of four keyboard keys (D, F, J, and K). Stimuli were either color words same as the ink color (congruent, n = 108) or different (incongruent, n = 108), or a string of letters “XXXXX” (neutral, n= 72). Feedback informed them whether they obtained a reward (reward, “$50c” or no reward, “$0c”) and whether the reward they received depended on their performance (performance-based feedback, a button graphic), or not (random feedback, a dice graphic). In order to earn rewards in the performance-based case, participants had to be both accurate and respond within an individually calibrated response deadline (see details below). The order of the two types of feedback was pseudo-randomized with half of the trials showing reward feedback first and the other half efficacy feedback. Every 2-4 trials the feedback was followed by a probe of efficacy (“How much do you think your rewards currently depend on your performance?”) or reward rate (“How often do you think you are currently being rewarded?”) to which participants responded on a visual analog scale ranging from 0 to 100. The number and timing of the probes was randomized per subject resulting in a median of 45 efficacy probes (SD=3.38) and 47 reward probes (SD=2.81).

**Figure 1.**
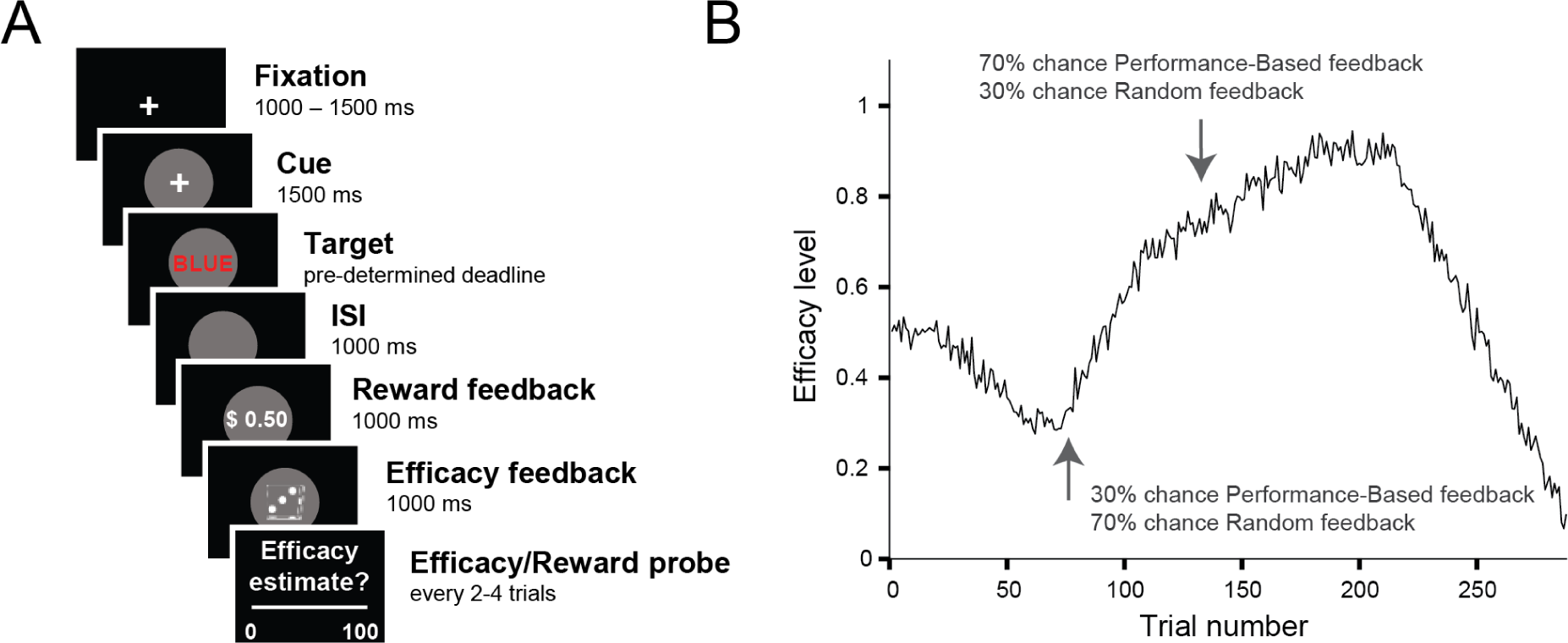
Manipulating expected efficacy and assessing learning in Study 1. A. Trial Schematic. On each trial participants saw a cue (gray circle), predicting the onset of a Stroop stimulus (target), and were then sequentially presented with reward and efficacy feedback. On half of the trials, efficacy feedback was presented first, and on the other half reward feedback was presented first. Every 2-4 trials participants were subsequently asked to estimate their current efficacy level (“How much do you think your rewards currently depend on your performance?”) or reward rate (“How often do you think you are currently being rewarded?”). **B. Efficacy manipulation.** We let the probability of performance-based vs. random feedback continuously drift over the course of the experiment (inversed for one half of the sample). Arrows mark time points with low and high efficacy, respectively. When efficacy was low, rewards were more likely to be random, whereas when efficacy was high, rewards were more likely to be performance-based.

Efficacy (performance-based or random rewards) on each trial was sampled from a binomial distribution with probabilities ranging between 0.1 and 0.9 that drifted over the course of the experiment and were predetermined (Figure 1B). In order to ensure that the performance-based and random trials did not differ in reward rate, reward feedback for the random trials was sampled from the moving window of the reward feedback of the previous 10 performance-based trials. At the beginning of the experiment a window with 8 rewards and 2 no rewards was used to reflect the pre-calibrated reward rate (details below), and this moving window was then updated after every trial. Thus, reward rate was not experimentally manipulated in the experiment and remained constant. We confirmed that reward rate was matched across performance-based and random trials by comparing reward probability between these trial types (*b =* 0.01; 95% CrI [- 0.07, 0.10]; p*_b_* _> 0_ = 0.38).

Prior to the main task, participants performed several practice phases of the Stroop task (approximately 15 minutes). First, they practiced the mappings between colors and keyboard keys (80 trials). Then they completed a short practice of the Stroop task with written feedback (“correct” or “incorrect”) on each trial (30 trials). Participants then completed 100 more of such trials during which we individually calibrated the reaction time deadline such that participants yielded approximately 80% reward rate. The reaction time calibration started from a fixed deadline of 750ms and increased or decreased this threshold in order to ensure that participants earn rewards on 80% of trials (i.e., that they are both accurate and below the deadline). The deadline obtained in this way (M = 796ms; SD = 73ms) was used in the main experiment and was not further adjusted. In the final practice phase participants were introduced to the two types of feedback which they would see in the main experiment (30 trials).

The experimental task was implemented in Psychophysics Toolbox (Brainard 1997; Pelli 1997; Kleiner et al. 2007) for Matlab (MathWorks Inc.) and presented on a 23 inch screen with a 1920 x 1080 resolution. All of the stimuli were presented centrally while the participants were seated 80 cm away from the screen.

#### Psychophysiological recording and preprocessing

EEG data were recorded at a sampling rate of 500 Hz from 64 Ag/AgCl electrodes mounted in an electrode cap (ECI Inc.), referenced against Cz, using Brain Vision Recorder (Brain Products, München, Germany). Vertical and horizontal ocular activity was recorded from below both eyes (IO1, IO2) and the outer canthi (LO1, LO2), respectively. Impedances were kept below 10 kΩ. Offline, data were processed using custom made Matlab scripts (Frömer et al. 2018) employing EEGlab functions (Delorme and Makeig 2004). Data were re-referenced to average reference, ocular artifacts were corrected using brain electric source analyses (Ille et al. 2002) based on separately recorded prototypical eye movements. The cleaned continuous EEG was then low pass filtered at 40 Hz and segmented into epochs around cue onset (-200 to 1500 ms), stimulus onset, and both efficacy and reward feedback (-200 to 800 ms). Baselines were corrected to the average of each 200 ms pre-stimulus interval. Segments containing artifacts, values exceeding ± 150 µV or gradients larger than 50 µV, were excluded from further analyses.

Single trial ERPs were then exported for further analyses in R (R Core Team 2017). The late CNV was quantified between 1000 and 1500 ms post neutral cue onset (Schevernels et al. 2014b; Frömer et al. 2016; Frömer et al. 2021a) as the average activity over 9 fronto-central electrodes (Fz, F1, F2, FCz, FC1, FC2, Cz, C1, and C2). The P3b was quantified between 350 and 500 ms (Fischer and Ullsperger 2013) for both reward and efficacy feedback and calculated as the averaged activity over 9 centro-parietal electrodes (Pz, P1, P2, POz, PO1, PO2, CPz, CP1, and CP2).

#### Learning models and statistical analyses

##### Learning models

Participants provided their subjective estimates of efficacy and reward every 4-8 trials (a total of 45 estimates), and we sought to fit a learning model to these estimates to be able to predict trial-by-trial adjustments in performance and neural markers of learning and cognitive control allocation. In order to obtain trial-by-trial estimates of efficacy and reward rate, we fitted two temporal difference learning models (Gläscher et al., 2010; Sutton & Barto, 2018) to the continuous subjective estimates of efficacy and reward rate (Rutledge et al., 2014; Eldar et al., 2016; Nagase et al., 2018). The first model (“1 learning rate efficacy model”) assumed that the estimate of efficacy for the next trial (*E*_*t*+1_) depended on the current efficacy estimate (*E*_*t*_) and the prediction error (δ_*t*_) weighted by a constant learning rate (α):

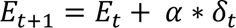

Where 0 ≤ *α* ≤ 1, and the prediction error is calculated as the difference between the contingency feedback on the current trial (*e*_*t*_) and the efficacy estimate on that trial: δ_*t*_ = *e*_*t*_ − *E*_*t*_. The model started from an initial value (free parameter) and updated the model-based efficacy estimate based on the binary efficacy feedback on each trial. For example, assuming a learning rate of 0.5 and the initial value of 0.5, the model would update the initial estimate following efficacy feedback signaling “performance-based” (*e*_*t*_ = 1) to 0.75. If on the next trial contingency feedback was “random” (*e*_*t*+1_ = 0) the model-based efficacy estimate would drop to 0.6. The model was fitted separately to the subjective estimates of efficacy with only the learning rate as a free parameter. The second model (“2 learning rates efficacy model”) was the same as the first model, but it included two learning rates: one learning rate for learning from the “performance-based” feedback, and another for learning from the “random” feedback. Finally, as a baseline, we also included the “intercept model” which did not update the efficacy estimate throughout the experiment, but just assumed that the estimate took one constant value. Importantly, the same models were fitted to obtain the model-based estimates of reward on each trial (“1 learning rate reward model” and the “2 learning rate reward model”). These models were fitted using trial-by-trial reward feedback and the subjective estimates of reward. The models were fit hierarchically to the data using maximum likelihood estimation (using mfit (https://github.com/sjgershm/mfit).

To calculate the likelihood of each data point, model-based estimates (0-1 range) were compared to the subjective efficacy estimates (range normalized to 0-1 range for each participant).

Likelihood was evaluated on trials which included a subjective estimate, as the likelihood that the difference between the model-based and the empirical estimate comes from a Gaussian distribution centered on 0 with a variance which was fitted as a free parameter each subject. This variance parameter served as the noise in the estimates. Likelihoods were log transformed, summed, and then maximized using the fmincon function in MATLAB.

We performed a parameter recovery study to show that the most complex model (the 2 learning rates model) can be successfully recovered. We simulated a dataset with the same number of trials and subjective efficacy or reward probes as in the actual experiment. We used the efficacy drifts presented to the actual subjects (half of the simulated subjects saw one drift, and half its inverse), and we used the reward feedback sequences of two actual subjects from our experiment. We simulated 200 agents which learned both efficacy and reward with the noise parameter fixed to 0.2, intercept fixed to 0.5, and the positive and negative learning rates sampled from a uniform distribution ranging from 0.001 to 0.5. These parameters were matched based on the range of values obtained from the empirical fits to our data. As shown in Figure S1 we were able to very reliably recover the simulated parameters for both efficacy and reward rate learning.

##### Statistical analyses

The efficacy and reward rate estimates obtained through fitting the learning model were then used to analyze the behavioral and EEG data. To this end, we fitted Bayesian multilevel regressions to predict subjective estimates of efficacy and reward rates, reaction times, accuracy, as well as the CNV and P3b amplitudes. Subjective estimates of efficacy and reward rate were regressed onto efficacy or reward feedback. Reaction times and accuracies were regressed onto trial-by-trial model-based estimates of efficacy and reward rate, as well as trial-by-trial CNV amplitude, while controlling for congruency. The P3b amplitudes were analyzed in two ways: with trial-by-trial model-based estimates of efficacy and reward rate and current feedback as predictors, and with model-based prediction errors and learning rates for each feedback type. CNV amplitudes were regressed onto trial-by-trial model-based estimates of efficacy. All of the fitted models controlled for the influence of the reward rate estimates.

Parallel analyses were done to predict the P3b in response to reward feedback, while controlling for the efficacy estimates.

The regression models were fitted in R with the *brms* package (Bürkner 2016) which relies on the probabilistic programming language *Stan* (Carpenter et al. 2016) to implement Markov Chain Monte Carlo (MCMC) algorithms and estimate posterior distributions of model parameters. The analyses were done based on the recommendations for Bayesian multilevel modeling using *brms* (Bürkner 2016; Bürkner 2017; Nalborczyk and Bürkner 2019). The fitted models included constant and varying effects (also known as fixed and random) with weakly informative priors (except for the behavioral and CNV analyses, see below for details) for the intercept and the slopes of fixed effects and the likelihoods appropriate for the modeled data (Ex-Gaussian for reaction times, Bernoulli for accuracy, and Gaussian for the subjective estimates and the EEG amplitudes). The fitted models included all of the fixed effects as varying effects.

All of the continuous predictors in the model were centered and the categorical predictors were contrast coded. Four MCMC simulations (“chains”; 20,000 iterations; 19,000 warmup) were run to estimate the parameters of each of the fitted models. The convergence of the models was confirmed by examining trace plots, autocorrelation, and variance between chains (Gelman and Rubin 1992). After convergence was confirmed, we analyzed the posterior distributions of the parameters of interest and summarized them by reporting the means of the distribution for the given parameter (*b*) and the 95% credible intervals (95% CrI) of the posterior distributions of that model. We report the proportion of the posterior samples on the relevant side of 0 (e.g., p*_b_* _< 0_ = 0.9), which represents the probability that the estimate is below or above 0. We also report Bayes factors (BF) calculated using the Savage-Dickey method (Wagenmakers et al. 2010). We report the BFs in support of the alternative hypothesis against the null (BF_10_), except for the analyses of accurate RT, accuracies, and CNV amplitude in which we have informative priors based on our previous study (Frömer et al. 2021a), and in which case we support the evidence in favor of the null (BF_01_).

To compare the positive and negative learning rates we fitted a model in which the learning rates were predicted by the learning rate type (Kruschke 2013). In this model we used Gaussian distributions (mean, standard deviation) as priors (intercept: (0.5,0.5); slopes: (0,0.5)).

We fitted two separate models to predict the subjective estimates of efficacy and reward rate based on previous feedbacks. At each timepoint the estimates were predicted by the current, and previous 4 feedbacks. The feedback on each of the trials (performance-based vs. random or reward vs. no reward) was entered as a constant effect and the models also included the intercept as a varying effect. As the subjective estimates could vary between 0 and 1, we used Gaussian distributions (intercept: (0.5,0.2); slopes: (0,0.2)) as priors.

For predicting the P3b amplitude in response to the onset of the efficacy feedback, we fitted two models. First, we fitted a model which included the model-based estimate of efficacy (prior to observing the current feedback), the actual feedback, and the interaction between the expected efficacy and the observed efficacy feedback. Additionally, we controlled for the reward rate estimate. Second, we fitted separate models which included the model-based prediction errors, the influence of the between-subject learning rates, and their interaction with the prediction errors, while controlling for the estimate of the reward rate. In this analysis the learning rates (one for feedback type for each subject) were mean-centered within subjects and thus any effect of the learning rates is driven by the difference between the random and the performance-based learning rate. For these models we selected wide Gaussian priors (intercept: (5,3); slopes (0,3)). The same logic in building models was applied for the analyses of the reward feedback. In these analyses we focused on the reward feedback processing and how it interacted with the model-based estimates of reward rates, while controlling for the model-based estimates of efficacy. We analyzed only the trials with correct responses for both the efficacy and the reward feedback analyses.

To test the influence of efficacy on the late CNV, we fitted a model which predicted the CNV based on the model-based efficacy estimates, while controlling for the effect of the reward rate estimates. Drawing on the results of our previous study (Frömer et al. 2021a), this model included Gaussian priors for the intercept (-0.16, 1.59) and the efficacy (-0.30, 0.73) and reward (0,0.73) slopes.

For predicting reaction times and accuracy we fitted models which included congruency (Facilitation: difference between neutral and congruent trials; Interference: difference between incongruent and neutral trials) and the model-based efficacy estimates, while controlling for the reward rate estimates. We used Gaussian distributions as informative priors based on our previous study (Frömer et al. 2021a), for both the reaction times (intercept: (624, 58.69); facilitation (15.54, 21.93), interference (61.12, 37.49); efficacy (-10.26, 19.51); reward (0, 19.51)) and accuracy_1_ analyses (intercept: (2.11, 0.81); facilitation (-0.45, 0.64), interference (-0.53, 0.81); efficacy (0.09, 0.32); reward (0, 0.32)).

To investigate how the late CNV influences the behavior, we fitted two models in which we predicted the reaction times and accuracy based on the CNV amplitude. The prior distributions for these models were weakly informative Gaussian distributions for predicting both the reaction times (intercept: (650, 200); slope: (0, 50)) and accuracy (intercept: (0.7, 0.2); slope: (0, 0.2)).

To visualize the topographies of the relevant ERP effects, we fitted the relevant models to all 64 channels and then plotted the posterior estimates of the effects of interest at each electrode (cf. Frömer et al., 2021b).

### Study 2

#### Participants

We recruited eighty-seven participants residing in the United States from Prolific – an online platform for data collection. Participants had normal or corrected-to-normal vision and gave informed consents. They were compensated with a fixed payoff of $8 per hour (median completion time of 74 minutes) plus a monetary bonus based on points earned during the task ($1 on average). The research protocol was approved by Brown University’s Institutional Review Board.

We a priori excluded participants who did not pass attention checks (N=8) or who took substantially longer than the average participant to complete the study (N=2 participants who took over 130 minutes), suggesting that they did not sustain attention to the experiment over that time. We fit our learning models to data from the remaining 77 subjects, and then excluded participants whose performance suggested inattention to the overall task (based on accuracies less than 70% across all trials – including the trials in which performance efficacy was low, N=6) or inattention to the task feedbacks and efficacy probes (based on low learning rates (N=19), and one subject with no variance in responses to reward probes). To identify participants with exceedingly low learning rates, we submitted all positive and negative efficacy learning rates to unsupervised Gaussian mixture models (as implemented in the Mclust package; (Scrucca et al., 2016) to determine the best fitting number and shape of clusters (model comparison via BIC).

This procedure identified four clusters of subjects with different overall learning rates (Figure S2B-C) and we excluded subjects from the first cluster as they all had very low learning rates relative to the other participants (both learning rates < 0.03). The subjects excluded based on low learning rates were most likely not paying attention to efficacy feedback, or were always giving very similar responses to the efficacy probes (Figure S3). The final sample included 51 participants (31 females, 20 males; median age = 29).

#### Experimental design

In order to better understand the computational mechanisms that lead to improved behavioral performance in high efficacy states (Study 1), we wanted to fit a Drift Diffusion model (DDM; Ratcliff & McKoon, 2008) to our behavioral data. If people allocate more attention when they think they have high performance efficacy, this should be observed as an increase in the drift rate (speed of evidence accumulation). However, Study 1 included a tight respond deadline for earning a reward, making it more challenging to fit the DDM. To avoid this issue, in Study 2 we adjusted the task to a free response paradigm which allowed us to investigate drift rate and threshold adjustments (cf. Leng et al., 2021). We used this design to test the hypothesis that higher efficacy estimates should predict increased drift rates.

Instead of single trials, participants now completed 288 intervals during which they could respond to as many trials (congruent and incongruent, removing the neutral condition) as they wished within a fixed time window (randomly selected between 2000, 3000, or 4000ms). Apart from this, the structure of the task remained the same: participants saw a fixation cross (1000, 1500, or 2000ms), then completed as many trials as they wished during a fixed interval, followed by the feedback (1500 ms) on how many points they earned and whether this was based on their performance or awarded to them based on random chance. Note that participants now received continuous reward feedback (10 points per correct response instead of the binary reward-no reward in Study 1). For example, if participants completed 4 trials correctly and 2 trials incorrectly within a performance-based interval they would receive 40 points and see feedback informing them that the points were based on their performance. To determine the number of points on random intervals, the same yoking procedure as in Study 1 was employed ensuring that the amount of reward was matched between performance-based and random intervals (reward amounts on random intervals were sampled from the moving average window of the past 10 performance-based intervals). We confirmed that the yoking procedure was successful by comparing the reward amounts on the two interval types (*b =* 0.00; 95% CrI [-0.00, 0.02]; p*_b_* _> 0_ = 0.16). As in Study 1, participants were probed every 2-4 intervals to estimate either how much they thought their rewards depended on their performance, or how often they were rewarded. We again implemented an efficacy drift (modified, but comparable to the drift in Study 1; Figure S2A), now across the 288 intervals of the task.

We gamified the task in order to make it more appealing for the participants. Instead of the Stroop task, we used a picture-word interference task in which four grey-scaled images of fruit (apple, pear, lemon, and peach) were overlaid by those fruit words. Participants responded to the image, while ignoring the word, by pressing one of the 4 corresponding keys. They first practiced this task, and then were introduced with a cover story telling them that they are in the garden and need to water the fruit in little patches by pressing the correct keys. They were instructed that they will be moving through a garden and that in some patches watering will directly translate into how many points they will be earning, while in the others that will not be the case (the efficacy drift). The experiment was implemented in Psiturk (Gureckis et al., 2016) and the participants performed the task on their own computers and were required to have a keyboard.

#### Learning models, Drift Diffusion model, and statistical analyses

##### Learning models

We fitted the same set of learning models as in Study 1, performed model comparison, and got the interval-by-interval model-based estimates of performance efficacy and reward. Note that in this version of the task participants earned points in each interval, unlike the binary rewards (reward vs. no reward) in Study 1. This meant that the reward learning model learned reward magnitudes rather than reward probabilities. However, model fitting and the further analyses were the same as in Study 1.

##### Statistical analyses

We fitted the same Bayesian multilevel models as in Study 1 to predict the influence of previous efficacy feedbacks on the subjective efficacy estimates, as well as the influence of the model-based efficacy estimates on reaction times and accuracies. For the analyses of the subjective efficacy estimates we used the same priors as in Study 1. For the analyses of the reaction times and accuracies, we used the posterior distributions obtained in Study 1 as the informative priors for the congruency, efficacy, and reward effects. The reaction times and accuracy models also controlled for the effects of the average congruency level in the interval and the interval length.

##### Drift Diffusion model

This model decomposes participant’s behavior into drift rate (the speed of evidence accumulation) and response threshold (the level of caution), allowing us to investigate which of these two components is affected by the efficacy estimates. We fitted the model using Bayesian hierarchical estimation as implemented in the HDDM package (Wiecki, Sofer, & Frank, 2013). The fitted model included the main effects of efficacy and reward rate estimates onto both drift rate and threshold, and included the effect of congruency on the drift rate. The responses were coded as correct or incorrect, and trials with reaction times below 250ms were excluded. All of the effects were allowed to vary across subjects, and we ran five MCMC chains (12,000 iterations; 10,000 warmup). We confirmed convergence by examining trace plots and variance between chains.

## Results

### Study 1

To investigate how efficacy estimates are learned, and how they affect control allocation, in Study 1 we recorded EEG while 40 participants performed a modified version of the Stroop task (Figure 1A). Across trials we varied whether reward outcomes ($0.50 vs. $0.00) were determined by a participant’s performance on a given trial (responding accurately and below a pre-determined response time criterion; *performance-based* trials) or whether those outcomes were determined independent of performance (based on a weighted coin flip; *random* trials).

Over the course of the session, we gradually varied the likelihood that a given trial would be performance-based or random such that, at some points in the experiment, most trials were performance-based (high efficacy level), and at other points most trials had random outcomes (low efficacy level) (Figure 1B). Importantly, unlike in our previous study (Frömer et al., 2021a), participants were not told whether a given trial would be performance-based (maximal efficacy) or random (minimal efficacy), but instead had to estimate their current efficacy level based on recent trial feedback, which indicated both the reward outcome ($0.50 vs. $0.00) and how that outcome was determined (performance-based [button symbol] vs. random [dice symbol]). We held reward rate constant across both feedback types by yoking reward rate on random trials to the reward rate on performance-based trials, and counter-balanced the time-course of the gradual change in efficacy (see Methods for details). To capture changes in efficacy expectations over the course of the session, we probed participants every 4-8 trials (averaging 44.3 probes per participant) to report their current estimates of efficacy. These efficacy probes were interleaved with probes asking participants to estimate the current reward rate, serving as foils and for control analyses.

### Participants dynamically update efficacy expectations based on feedback

To determine whether and how participants learned their current efficacy levels, we first analyzed the influence of previous efficacy feedback (whether outcomes on a given previous trial had been performance-based or random) on one’s current subjective estimates of efficacy. We found very strong evidence that participants adjusted their subjective efficacy upward or downward depending on whether the most recent trial was Performance-Based or Random (*b =* 0.14; 95% CrI [0.12, 0.16]; p*_b_* _< 0_ = 0; BF_10_ > 100), and that this remained true (but to diminishing degrees) up to five trials back (all p*_b_* _< 0_ < 0.01; Figure 2A; Table S1). This effect of feedback was present only for the efficacy feedback, while reward feedback did not predict the subjective estimates of efficacy (Figure 2A). These results suggest that participants were dynamically updating their efficacy estimates based on efficacy feedback.

**Figure 2.**
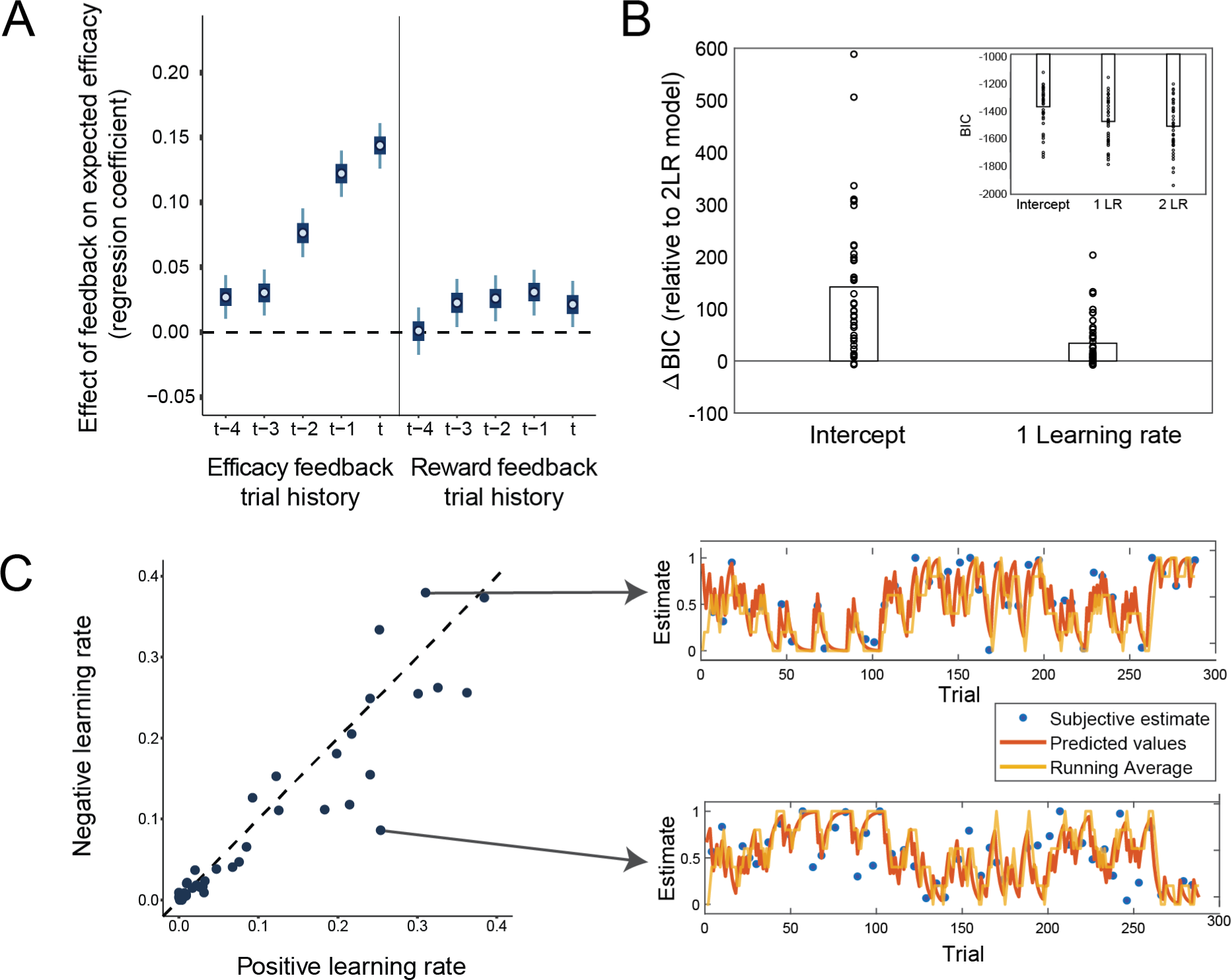
Efficacy learning is captured by a reinforcement learning model with separate learning rates for performance-based and random feedback in Study 1. A. Efficacy estimates track recent efficacy feedback. Regression weights for the influence of the current (t) and previous contingent vs. random feedback, as well as reward feedback, on the subjective efficacy estimate. Error bars represent 50% and 95% highest density intervals. B. A separate learning rate model captures efficacy learning best. BICs of fitted intercept only and one learning rate model relative to two learning rate models are plotted for each participant and favor the two learning rate model. C. Learning rate biases vary between participants. Positive and negative learning rate estimates are plotted for all participants. Points below the diagonal indicate higher learning rates for performance-based compared to random feedback, and points above the opposite. D. Example learning trajectories. Subjective and model-based efficacy estimates, and a running average of the previous 5 efficacy feedbacks, for a participant with a higher learning rate for the contingent compared to random feedback (upper) and a participant with the reverse bias (lower).

This pattern of learning was accounted for by a standard reinforcement learning (RL) algorithm, the delta-rule, according to which efficacy estimates are updated in proportion to the prediction error between the expected efficacy and the efficacy feedback on a given trial (i.e., whether a given outcome was determined by performance or not) (Sutton and Barto 2018).

Interestingly, consistent with studies of reward learning (Niv et al. 2012; Collins and Frank 2014; Lefebvre et al. 2017; Chambon et al. 2020; Garrett and Daw 2020), we found that the RL model that best accounted for our data allowed efficacy estimates to be updated differently from trials that were more efficacious than expected (Performance-Based feedback) than from trials that were less efficacious than expected (Random feedback), by having separate *learning rates* scaling prediction errors in the two cases. Even when accounting for model complexity, this Two Learning Rate Efficacy model outperformed a One Learning Rate Efficacy model as well as a baseline model that fits a single constant efficacy estimate and no learning (Intercept Model) (Figure 2B). In addition, we were able to successfully recover the parameters of this model from a simulated dataset matched to our data (see Methods and Figure S1). We found that the two learning rates for this best-fit model varied across the group (Figure 2E), but did not find that one was consistently higher than the other (*b =* 0.02; 95% CrI [-0.04, 0.08]; p*_b_* _< 0_ = 0.260; BF_01_=13.55; Figure S4), despite most participants (80%) tending to learn more from Performance-Based than Random trials. Finally, model-based efficacy estimates were strongly related to the raw subjective estimates on trials on which participants reported efficacy (*b =* 0.77; 95% CrI [0.62, 0.91]; p*_b_* _< 0_ < 0.001; BF_01_ > 100; R^2^=0.50), demonstrating that the model successfully captured the raw estimates. Taken together, these results show that participants dynamically updated their expected efficacy based on trial-by-trial feedback, and that they did so differentially based on whether the trial was more or less efficacious than expected. The fitted models further enable us to generate trial-by-trial estimates of expected efficacy and efficacy prediction errors, which we use in model-based analyses of behavior and neural activity below. Note that in all the following analyses we control for reward estimates obtained from models fit to reward feedback (for details see below).

### The feedback-related P3b indexes updating of efficacy expectations

To investigate the neural mechanism underlying feedback-based learning of efficacy, we probed the centroparietal P3b ERP component (Figure 3A), an established neural correlate of prediction-error based learning (Fischer and Ullsperger 2013; Nassar et al. 2019). If the P3b indexes feedback-based updating of *efficacy* predictions, as it does for reward predictions, we would expect this ERP to demonstrate several key markers of such a learning signal. First, we would expect the P3b to reflect the extent to which efficacy feedback (Performance-Based vs.

**Figure 3.**
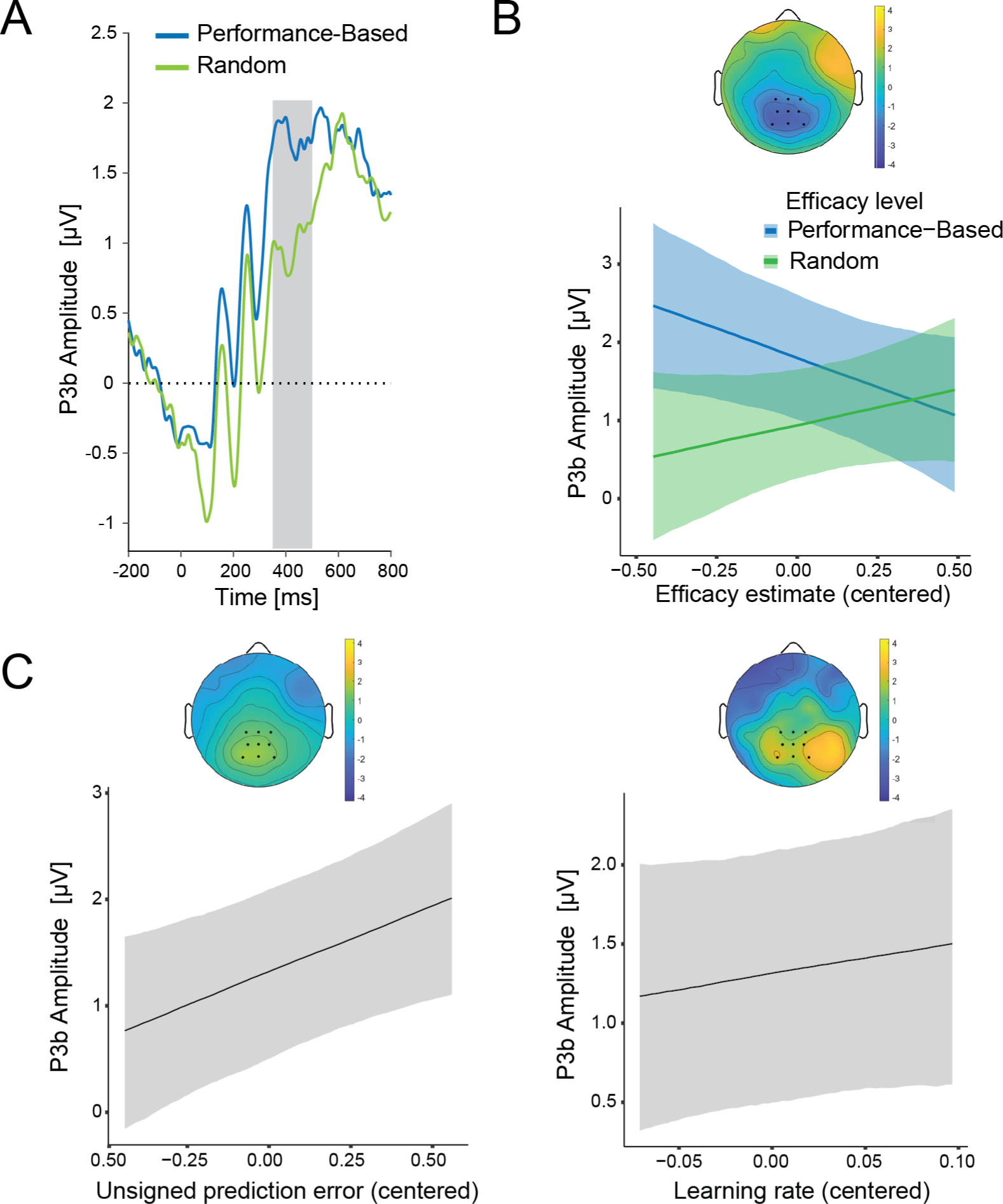
P3b reflects dynamically changing efficacy estimates during processing of efficacy feedback in Study 1. **A.** ERP average for the P3b locked to the onset of the efficacy feedback separately for performance-based and random feedback. The grey area shows the time window used for quantifying the P3b. **B.** LMM predicted P3b amplitudes are plotted for performance-based and random feedback as a function of efficacy estimates. The topography shows the interaction of efficacy estimate with efficacy feedback in the P3b time window. **C.** Predicted (centered) effects of unsigned prediction errors (left) and model-based learning rates (right) on the P3b. Shaded error bars represent 95% confidence intervals. Topographies display fixed-effects estimates.

Random) deviates from the current level of expected efficacy. In other words, the P3b should track the magnitude of the *unsigned efficacy prediction error (PE)* on a given trial. We tested this by examining how the amplitude of the P3b to a given type of efficacy feedback varied with model-based estimates of the participant’s efficacy expectation on that trial, while holding the expected reward rate constant (see the Methods and Materials sections for the details of the experimental design and the statistical models). If the P3b signaled efficacy PE then its amplitude should scale inversely with the expected probability of a given type of feedback (i.e., how *unexpected* that feedback is), thus correlating *negatively* with expected efficacy on trials providing Performance-Based feedback and correlating *positively* with expected efficacy on trials providing Random feedback. In addition to overall higher P3b to performance-based compared to random feedback (*b* = 0.86; 95% CrI [0.42, 1.31]; p*_b_*_<0_ = 0; BF_10_ =30.86; Figure 2B), we found exactly this predicted interaction between feedback type and expected efficacy (*b =* -2.40; 95% CrI [-4.07, -0.74]; p*_b_* _> 0_ = 0; BF_10_ = 24.40; Figure 2B; Figure S5A; Table S2), with the P3b amplitude decreasing with model-based estimates of expected efficacy on Performance- Based trials (b = -1.49; 95% CrI [-2.72, -0.27]; p*_b_* _> 0_ = 0.03; BF_10_ = 1.52) and increasing numerically, but not robustly with expected efficacy on Random trials (*b =* 0.91; 95% CrI [-0.34, 2.21]; p*_b_* _< 0_ = 0.12; BF_10_ = 0.44). Accordingly, when we regressed P3b amplitude on our model- based estimates of trial-to-trial efficacy PE, we found a positive relationship (*b* = 1.25; 95% CrI [0.35, 2.15]; p*_b_*_<0_ = 0.01; BF_10_ = 5.84; Figure 3C-left; Table S2).

In addition to tracking efficacy PEs, the second key prediction for the P3b if it indexes efficacy learning, is that it should track the extent to which PEs are used to update estimates of efficacy (i.e., the learning rate). In the current study, we found that participants differed in their learning rates for the two forms of efficacy feedback (Performance-Based vs. Random), providing us with a unique opportunity to perform a within-subject test of whether P3b tracked differences in learning rate across these two conditions. Specifically, we could test whether a given subject’s P3b was greater in the feedback condition for which they demonstrated a higher learning rate. However, we have not found conclusive evidence for the increase in P3b for the within-subject feedback condition with the higher learning rate (*b* = 2.00; 95% CrI [-2.21, 6.04]; p*_b_*_<0_ = 0.17; BF = 1.08; Figure 3C-right). While, theoretically, prediction error and learning rate would interact in predicting the P3b amplitude, we did not observe such an interaction here. This finding is in line with previous work on reward processing (Fischer and Ullsperger 2013), which has found additive effects of prediction errors and learning rate on P3b. While we found the effect of prediction errors, our learning rate effect was going in the expected direction, but was not conclusive.

### The CNV indexes control allocation based on updated expectations of efficacy

Thus far, our findings suggest that participants dynamically updated expectations of their performance efficacy based on feedback, and that the P3b played a role in prediction error-based updating of these expectations. Next, we tested the prediction that learned efficacy estimates determine the expected benefits of control, and thus influence how much control is allocated (Shenhav et al. 2013). We have previously shown that people exert more control when expecting their performance to be more rather than less efficacious on the upcoming trial (Frömer et al., 2021a). This was reflected in better behavioral performance and higher amplitudes of the contingent negative variation (CNV; Figure 4B-left) - a slow negative fronto-central waveform preceding target onset, which is increasingly negative as the amount of control allocated in preparation for the task increases (Grent-’t-Jong and Woldorff 2007; Schevernels et al. 2014a). Here we used the same marker of control, but, unlike in our previous study, efficacy expectations were (a) learned rather than explicitly instructed; (b) varied over time rather than having a fixed association with a cue; and (c) varied continuously across the range of efficacy probabilities rather than alternating between maximal and minimal efficacy. We were therefore interested in testing whether these dynamically varying expectations of efficacy, as estimated by our model, would still exert the same influence on behavior and neural activity.

**Figure 4.**
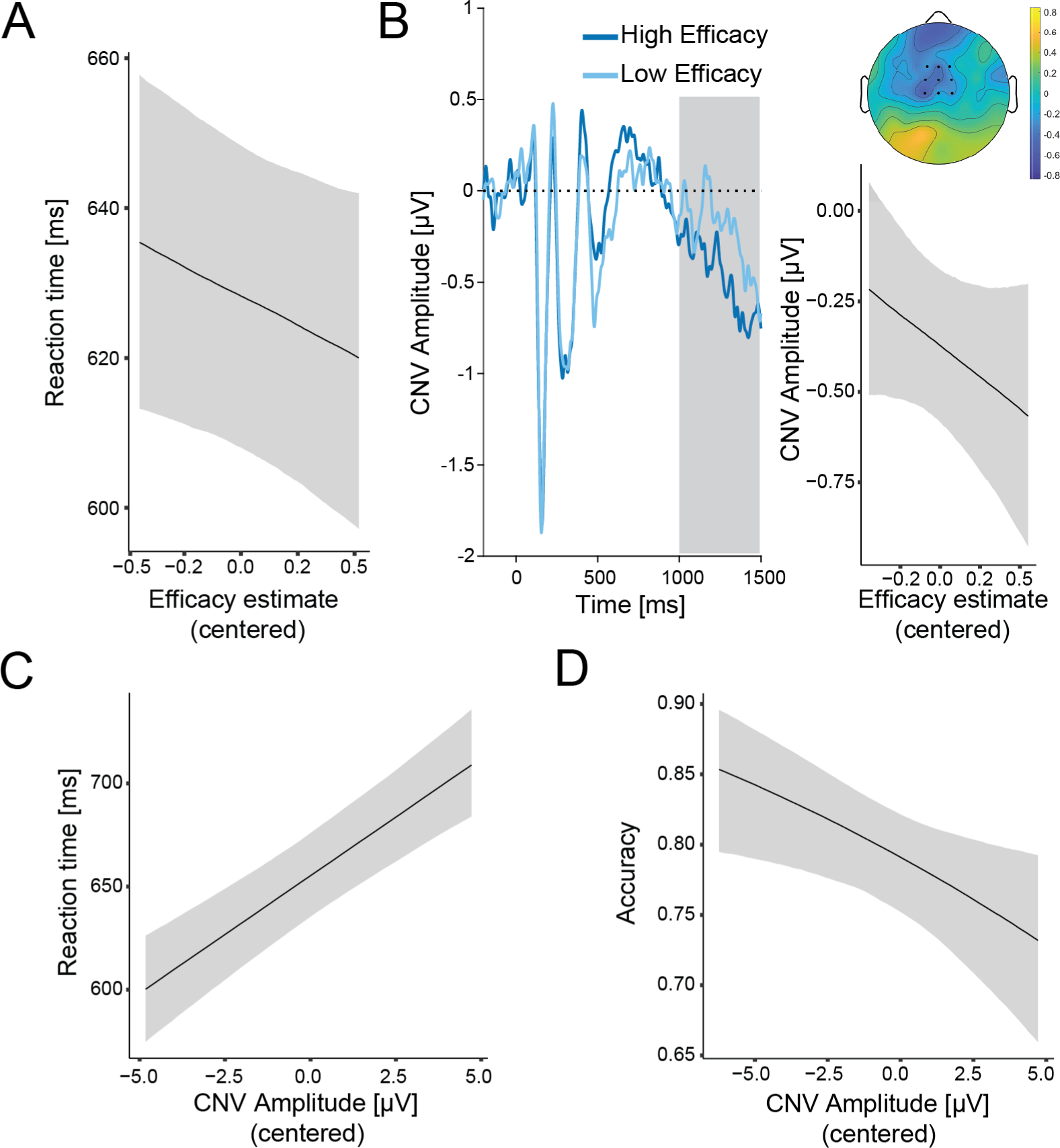
Efficacy estimates influence allocation of control and behavior in Study 1. **A.** Higher efficacy predicts faster accurate responses. **B. CNV increases with higher efficacy.** Left: Grand-average ERP for high and low efficacy estimates (median split used for plotting). The shaded area marks the time window used for CNV quantification. Time 0 corresponds to the onset of the neutral cue. Right: Predicted CNV amplitudes as a function of efficacy estimates. The topography shows the fixed effect of the efficacy estimate from the fitted linear model. **C. -D. Larger CNV amplitude predicts better performance.** Predicted accurate RT (**C.**) and accuracy **(D.)** as a function of efficacy estimates. Shaded error bars indicate 95% confidence intervals.

Consistent with our predictions and our previous findings, participants tended to perform better when they expected performance to be more efficacious, responding faster on correct trials with increasing model-based estimates of efficacy (Figure 4A; Figure S6A; Table S3). This finding replicates the performance effects we observed using instructed cues, albeit with only modest evidence (*b =* -16.00; 95% CrI [-34.91, 2.96]; p*_b_* _> 0_ = 0.05; BF_01_ = 1.68). Like in our previous studies, faster responding was not explained by a change in speed-accuracy trade-off, as accuracy did not decrease with increasing efficacy (*b =* 0.12; 95% CrI [-0.20, 0.44]; p*_b_* _> 0_ = 0.23; BF_01_ = 1.99; Figure S6A). These analyses controlled for the standard behavioral effects related to Stroop congruency (i.e., slower and less accurate responding for incongruent relative to congruent trials; Figure S7), as well as for the reward rate estimates.

If the CNV provides an index of control allocation based on current incentive expectations, it should both reflect one’s latest efficacy estimate and should predict performance on the upcoming trial. Our findings support both predictions. Regressing single-trial CNV amplitude onto model-based efficacy estimates, and controlling for expectations of reward (discussed below), we found that the CNV amplitudes had a positive relationship^2^ with the current efficacy expectations (*b =* -0.35; 95% CrI [-0.85, 0.16]; p*_b_* _> 0_ = 0.09; BF_10_ = 2.73; Figure 4B-right; Figure S5B; Table S4). However, this effect was weaker than in the previous experiment with cued efficacy levels, which is to be expected given that in this experiment participants had to learn their efficacy levels. As with the behavioral finding above, this result provides evidence consistent with our previous CNV finding using instructed cues. We further replicate our earlier finding (Frömer et al., 2021a) that larger CNV amplitude in turn predicted better performance on the upcoming trial, with participants responding faster (*b =* 11.41; 95% CrI [8.09, 14.72]; p*_b_* _< 0_ = 0; BF_10_ > 100; Figure 4C; Table S5) and more accurately (*b =* -0.07; 95% CrI [-0.12, -0.01]; p*_b_* _> 0_ = 0.01; BF_10_ = 3.14; Figure 4D; Table S5) as CNV increased.

### Parallel learning of efficacy and reward rate

We held the amount of reward at stake constant over the course of the experiment, but the frequency of reward (reward rate) varied over the course of the session based on a participant’s recent performance, and participants were sporadically probed for estimates of their reward rate (interleaved with trials that were followed by efficacy probes). Our efficacy manipulation explicitly controlled for this variability by yoking Random-Outcome feedback to a participant’s recent reward rate (see methods for details). However, this additional source of variability also provided an opportunity to examine additional mechanisms of learning and adaptation in our task. As in the case of efficacy estimates, reward rate estimates were robustly predicted by reward feedback on previous trials (Table S1; Figure S8A), and this reward rate learning process was well captured by a two learning rate reward rate model (Garrett and Daw 2020; S8B-C), with the model-based estimates successfully predicting the reported subjective estimates (*b =* 0.82; 95% CrI[0.60, 1.02]; p*_b_* _< 0_ < 0.001; BF_01_ > 100; R^2^=0.58; Figure S8B-C). Updates to these reward rate estimates were reflected in P3b modulations of (unsigned) reward prediction errors and associated learning rates (Fischer and Ullsperger 2013; Figure S9; Table S6). This pattern of results provides additional evidence that efficacy learning involves similar neural and computational mechanisms as reward-based learning.

### Study 2

#### Learned efficacy modulates control over information processing

Our findings suggest that people rely on domain-general mechanisms to learn about their performance efficacy in a given environment, and use these learned estimates of efficacy to optimize performance on their task. Specifically, in Study 1 we found that higher levels of learned efficacy are associated with faster responses, albeit with modest evidence (*b =* -16.00; 95% CrI [-34.91, 2.96]; p*_b_* _> 0_ = 0.05; BF_01_ = 1.68). We also found that this speeding occurred on correct but not incorrect trials, suggesting that these performance adjustments reflected attentional control rather than adjustments to speed-accuracy tradeoffs. However, these findings remain only suggestive in the absence of a formal model, and the presence of a stringent response deadline in this study (individually calibrated for each subject during the practice phase to ensure the reward rate of 80%; M = 796ms; SD = 73ms) presented an obstacle to fitting our behavioral data to such a model without additional assumptions (e.g., regarding the form of a collapsing threshold).

To provide further support for our proposal that learned efficacy influences control over information processing, we ran an additional behavioral study (Study 2). Participants in this study (N = 51) performed a web-based version of the task in Study 1, with the biggest modification being that the Stroop trials (now using picture-word rather than color-word interference) were performed over the course of short self-paced time intervals rather than trial by trial as in Study 1. Specifically, participants were given limited time windows (2-4s) to complete as many Stroop trials as they wanted to and were rewarded at the end of each interval (cf. Leng et al., 2021). When rewards were performance-based, participants received a number of points exactly proportional to the number of correct responses they gave during that window.

When rewards were random, the number of points was unrelated to performance on that interval but (as in Study 1) was yoked to their performance in previous performance-contingent intervals. Following our approach in Study 1, we varied the likelihood of a given interval being performance-based or random over the course of the session (Figure S2A), and sporadically probed participants for their subjective estimates of expected efficacy and reward rate. While in most respects very similar to the paradigm in Study 1, one noteworthy feature of this self-paced design is that it resulted in a much less stringent deadline for responding, thus producing reaction time patterns more typical of free-response paradigms for which the traditional (fixed-threshold) DDM was designed. Note that because this was an online sample we also employed additional cutoff criteria to exclude inattentive participants, as detailed in the Methods section and Figures S2B-C and S3.

Replicating the learning patterns observed in Study 1, subjective estimates of efficacy in Study 2 reflected a running average over recent efficacy feedback (*b =* 0.18; 95% CrI [0.16, 0.20]; p*_b_* _< 0_ = 0; BF_10_ > 100). This effect was again weighted towards the most recent feedback but still present up to five intervals back (all p*_b_* _< 0_ < 0.01; Figure 5A, Figure S10, and Table S7). As in Study 1, this learning pattern was best captured by an RL algorithm with two learning rates (Figure 5B), though positive and negative efficacy learning rates did not significantly differ from one another on average (*b =* 0.03; 95% CrI [-0.03, 0.08]; p*_b_* _< 0_ = 0.260; BF_01_=11.74; Figure 5C). As in Study 1, the model-based efficacy estimates successfully predicted the raw subjective estimates on intervals on which participants reported their efficacy beliefs (*b =* 0.86; 95% CrI [0.81, 0.90]; p*_b_* _< 0_ < 0.001; BF_01_ > 100; R_2_ = 0.60), and the same was true for the model-based reward estimates predicting the subjective reward estimates (*b =* 1.00; 95% CrI[0.95, 1.04]; p*_b_* _< 0_ < 0.001; BF_01_ > 100; R^2^=0.66).

**Figure 5.**
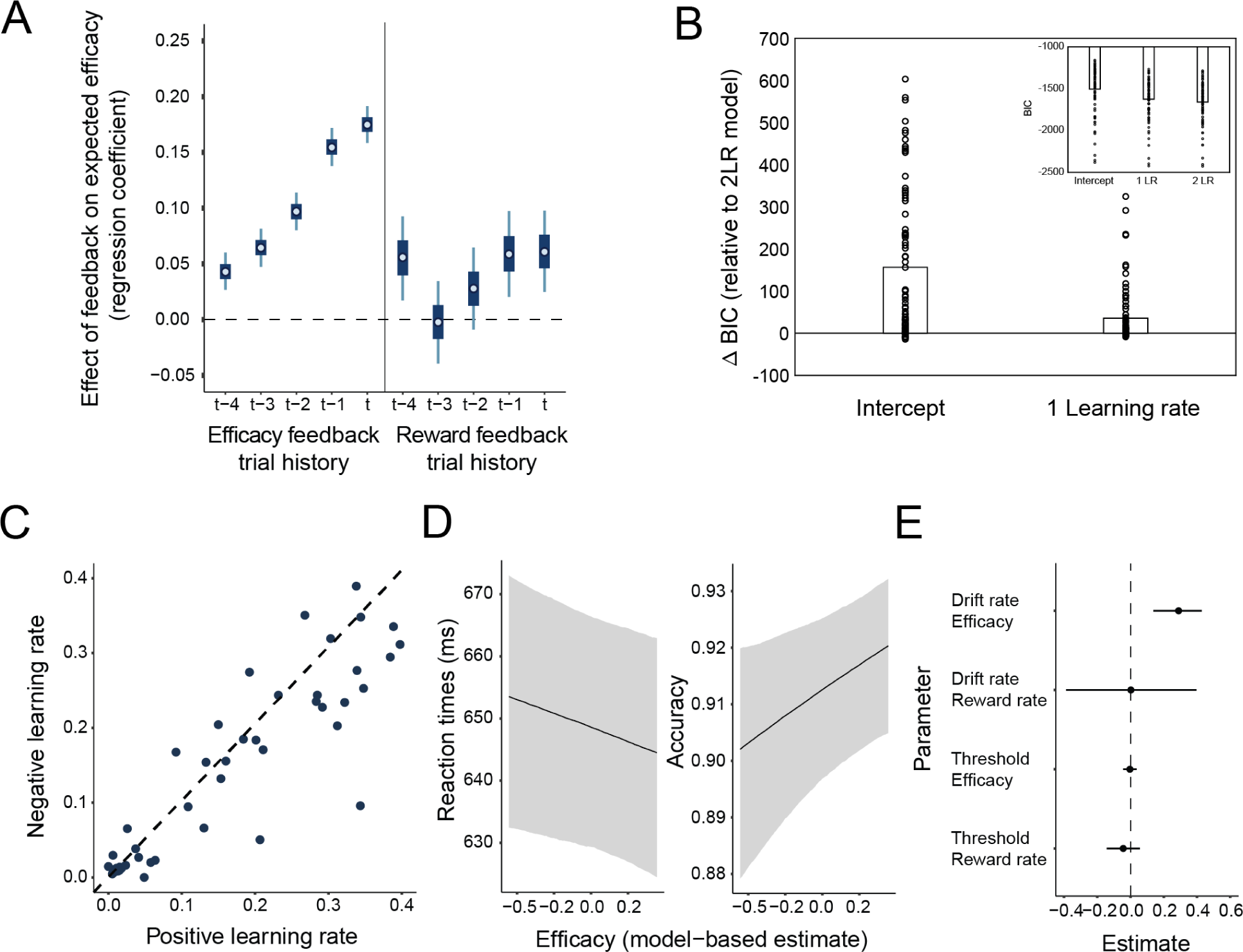
Efficacy is learned in the same way in a modified task and influences a behavioral marker of control allocation. **A.** Regression weights for the influence of previous feedback type (efficacy and reward) on the subjective efficacy estimate. Error bars represent 50% and 95% highest density intervals. **B. Two learning rate model captures efficacy learning best.** Differences in BICs between the two learning rate model, and the intercept-only and one learning rate models. **C. Efficacy learning rates.** Positive and negative efficacy learning rates for each participant**. D. Higher model-based efficacy estimates predict better behavioral performance.** Higher efficacy estimates reduce reaction times (left) and improve accuracy (right). **E. Higher model-based efficacy estimates predict increased allocation of attention.** Parameter estimates from the drift diffusion model. Higher efficacy estimates increase drift rates, but not response caution (thresholds).

Critically, we once again found that higher model-based estimates of efficacy predicted better performance on the upcoming interval. Participants responded faster (*b =* -10.25; 95% CrI [-19.86, -0.20]; p*_b_* _> 0_ = 0.02; BF_01_ = 2.33) and more accurately (*b =* 0.23; 95% CrI [0.03, 0.43]; p*_b_* _> 0_ = 0.01; BF_01_ > 100) with increasing model-based efficacy estimates (Figure 6B, Figure S6B, and Table S8). To formally test whether these behavioral patterns reflected adjustments in information processing (i.e., the rate of evidence accumulation once the stimuli appeared) or instead reflected adjustments in speed-accuracy tradeoffs (i.e., the threshold for responding), we fit these data with the hierarchical drift diffusion model (HDDM; Wiecki, Sofer, & Frank, 2013). We tested whether model-based efficacy estimates predicted trial-by-changes in drift rate, threshold, or both, while controlling for influences of expected reward rate on those same DDM parameters. We found that higher levels of expected efficacy were associated with higher drift rates (*b =* 0.07; 95% CrI [0.14, 0.43]; p*_b_* _< 0_ = 0.00) but were uncorrelated with threshold levels (*b =* -0.00; 95% CrI [-0.04, 0.04]; p*_b_* _< 0_ = 0.74). Expected reward rate was not correlated with either drift rate or threshold (Table S9). These results suggest that participants responded to changes in performance efficacy by adjusting their attention to the task, rather than simply adjusting their response threshold (i.e., becoming more or less impulsive).

## Discussion

To evaluate the expected benefits of investing cognitive control into a task, people need to consider whether this control is necessary and/or sufficient for achieving their desired outcome (i.e., whether these efforts are worthwhile). A critical determinant of the worthwhileness of control is performance efficacy, the extent to which performance on a control-demanding task matters for outcome attainment versus those outcomes being determined by performance-unrelated factors. However, the mechanisms through which people estimate the efficacy of their performance based on previous experience are largely unknown. Here, we identified the computational and neural mechanism through which efficacy is learned and used to guide the allocation of cognitive control. Across two experiments, we found that participants dynamically updated expectations of the efficacy of their task performance (i.e., the likelihood that this performance will determine reward attainment), and used those expectations to adjust how much control they allocated. The feedback-based updating of efficacy was well-captured by a prediction error-based learning algorithm. Model-based estimates of efficacy and efficacy prediction errors were encoded by canonical neural signatures of effort allocation and belief updating, respectively. Importantly, these findings cannot be explained by variability in reward, as reward rate was held constant across efficacy levels, and the subjective reward rate was controlled for statistically. Further, our model-based analysis revealed that people allocated more control when they learned that they had more efficacy, extending our previous findings on instructed efficacy (Frömer et al., 2021a). Taken together, our results uncover the mechanism through which efficacy estimates are learned and used to inform mental effort investment within a given task environment.

Previous research has characterized the learning algorithms people use to learn the reward value of different states and actions in their environment (Gläscher et al. 2010; Sutton and Barto 2018). Recent theoretical (Jiang et al. 2014; Lieder et al. 2018; Verbeke and Verguts 2019) and empirical (Bejjani et al. 2018; Otto and Daw 2019; Jiang et al. 2020; Bustamante et al. 2021 Jan 6) work has extended this research to show how similar algorithms guide learning and adaptation of cognitive control under varying rewards and task demands within a given task environment. Our findings extend this work further in several ways. First, we show that people leverage weighted prediction errors when learning about the efficacy of task performance, independently of potential reward and task difficulty. Second, we show that they update their efficacy expectations differently depending on whether performance was more efficacious or less efficacious than they expected, demonstrating a striking parallel with dual learning rate models that have been found to prevail in research on reward learning (Collins and Frank 2013; Lefebvre et al. 2017; Garrett and Daw 2020), including in our own reward rate data. Third, we show that participants dynamically adjust their control allocation based on learned efficacy, just as they do for learned rewards and task demands (Bugg et al. 2011; Jiang et al. 2014; Lieder et al. 2018).

Our neural findings build further on past research on learning and control adaptation. The P3b component has been shown to track prediction-error based learning from action-relevant outcome values (Fischer and Ullsperger 2013; Nassar et al. 2019; Lohse et al. 2020). Here we show that this neural marker tracks learning about efficacy in the same way as it tracks learning about rewards. We found increased P3b amplitudes when people experienced feedback about outcome contingencies that was less expected given their current estimate of efficacy (e.g., expecting low efficacy, but getting performance-contingent feedback), relative to when these contingencies were more expected (e.g., expecting low efficacy and getting random feedback). Our additional finding that P3b amplitude was overall larger for efficacy compared to no efficacy feedback demonstrates that our participants were not just tracking the frequency of the two types of feedback, as predicted by an oddball account. Instead this finding suggests that they were actively learning from the feedback.

Extending previous findings on cueing efficacy and/or reward (Schevernels et al. 2014b; Frömer et al. 2021a), our CNV and behavioral results further show that participants used these learned efficacy estimates to calibrate their effort and their performance. Notably, unlike in previous work, our study shows effort-related changes in CNV amplitude entirely divorced from perceptual cues, providing evidence that this activity truly reflects adjustments in control, rather than reactive processing of features associated with the cue. Taken together, our findings suggest that similar neural mechanisms underlie learning and control adaptation in response to variability in one’s efficacy in a task environment, as they do in response to variability in expected rewards (Leng et al., 2021; Otto & Daw, 2019). By fitting behavioral data from this task to a drift diffusion model (Study 2), we were able to further demonstrate that participants were adapting to changes in expected efficacy by enhancing the processing of stimuli (i.e., increasing their rate of evidence accumulation) - potentially via attentional control mechanisms – rather than by adjusting their threshold for responding. This particular pattern of control adjustments was predicted for the current task because performance-contingent rewards depended on responses being *both* fast and accurate (as in Frömer et al., 2021a), but future work should test the prediction that different control adjustments should emerge under different performance contingencies (cf. Leng et al., 2021; Ritz et al., 2021).

Our findings build on previous research on how people learn about controllability of their environment. Studies have examined the neural and computational mechanisms by which humans and other animals learn about the contingencies between an action and its associated outcome, and demonstrated that these learned action-outcomes contingencies influence which actions are selected and how vigorously they are enacted (Dickinson and Balleine 1995; Liljeholm et al. 2011; Manohar et al. 2017; Moscarello and Hartley 2017; Ly et al. 2019). Our work extends this research into the domain of cognitive control, where the contingencies between actions (i.e., control adjustments) and outcomes (e.g., reward) depend both on whether control adjustments predicts better performance and whether better performance predicts better outcomes (Shenhav et al., 2021). Learning control-outcome contingencies therefore requires learning about how control states map onto performance (control efficacy) as well as how performance maps onto outcomes (performance efficacy). By describing the mechanisms by which people solve the latter part of this learning problem, and demonstrating that these are comparable to those engaged during action-outcome learning, our study lays critical groundwork for better understanding the links between selection of actions and control states.

Our efficacy-updating results are a reminder that many aspects of feedback are reflected in prediction error signals (Langdon et al. 2018; Frömer et al. 2021b). In the present study, we intentionally separated feedback about reward and efficacy to isolate the cognitive and neural learning mechanisms associated with each. In doing so, we have taken an important first step towards understanding the updating mechanisms underlying each. Further work is needed to understand how they are inferred in more naturalistic settings, in which different forms of feedback are often multiplexed, containing information about the values of actions that were taken as well as about the features and structure of the environment (cf. Dorfman et al., 2019; 2021).

Another distinct element of more complex naturalistic environments is that the same feedback can be used to evaluate multiple targets, internal ones, such as the selected response and its predicted outcome, or external ones, such as the source of feedback/environment (Carlebach and Yeung 2020). Such multi-level prediction error signals might for instance explain, why despite close links between P3b and behavioral adaptation (Yeung and Sanfey 2004; Chase et al. 2011; Fischer and Ullsperger 2013), this link is context dependent and attempts to link the P3b uniquely to behavioral updating have failed (Nassar et al., 2019). Reinforcement learning, and predictive inference more generally, have been proposed to not only support the selection of individual actions, but also extended sequences of actions and control signals (Holroyd and Yeung 2012; Lieder et al. 2018). Alongside evaluations of actions and environmental states, neural signatures of feedback-based learning could thus further reflect evaluations of control signals, their quality or intensity. Given the many potential causes a given outcome can have, and the flexibility that people have in how they use the feedback, it is easy to see how feedback could be misattributed and lead to inaccurate beliefs about performance efficacy. Such beliefs about environmental statistics can drive changes in feedback-processing and behavioral adaptation, above and beyond the true statistics (Schiffer et al. 2017), and are thus of particular importance for understanding some of the cognitive symptoms of mental disorders.

Understanding how efficacy estimates develop based on previous experiences is crucial for understanding why people differ in how much cognitive control they allocate in different situations (Shenhav et al.). People differ in their beliefs about how much control they have over their environments (Leotti et al. 2010; Moscarello and Hartley 2017), and in their ability to estimate their efficacy (Cohen et al. 2020). Further, many mental disorders, including depression and schizophrenia, have been linked with one’s estimates of their ability to control potential outcomes in their environment, including through allocation of control (Huys and Dayan 2009; Maier and Seligman 2016), and we have recently proposed that such changes can drive impairments of motivation and control in those populations (Grahek et al. 2019). As we show in this study, when people have learned to expect low efficacy, they will allocate less cognitive control, which can manifest as apparent control deficits. The experimental and modeling approach taken in our study helps uncover a more fine-grained view of how components of motivation are learned and used to support the allocation of cognitive resources. In this way, our study takes a first step toward a better computational and neural account of efficacy learning, which can aid the understanding of individual differences in the willingness to exert mental effort, as well as the development of interventions aimed at teaching individuals when these efforts truly matter.

## Supplementary materials

**Figure S1.**
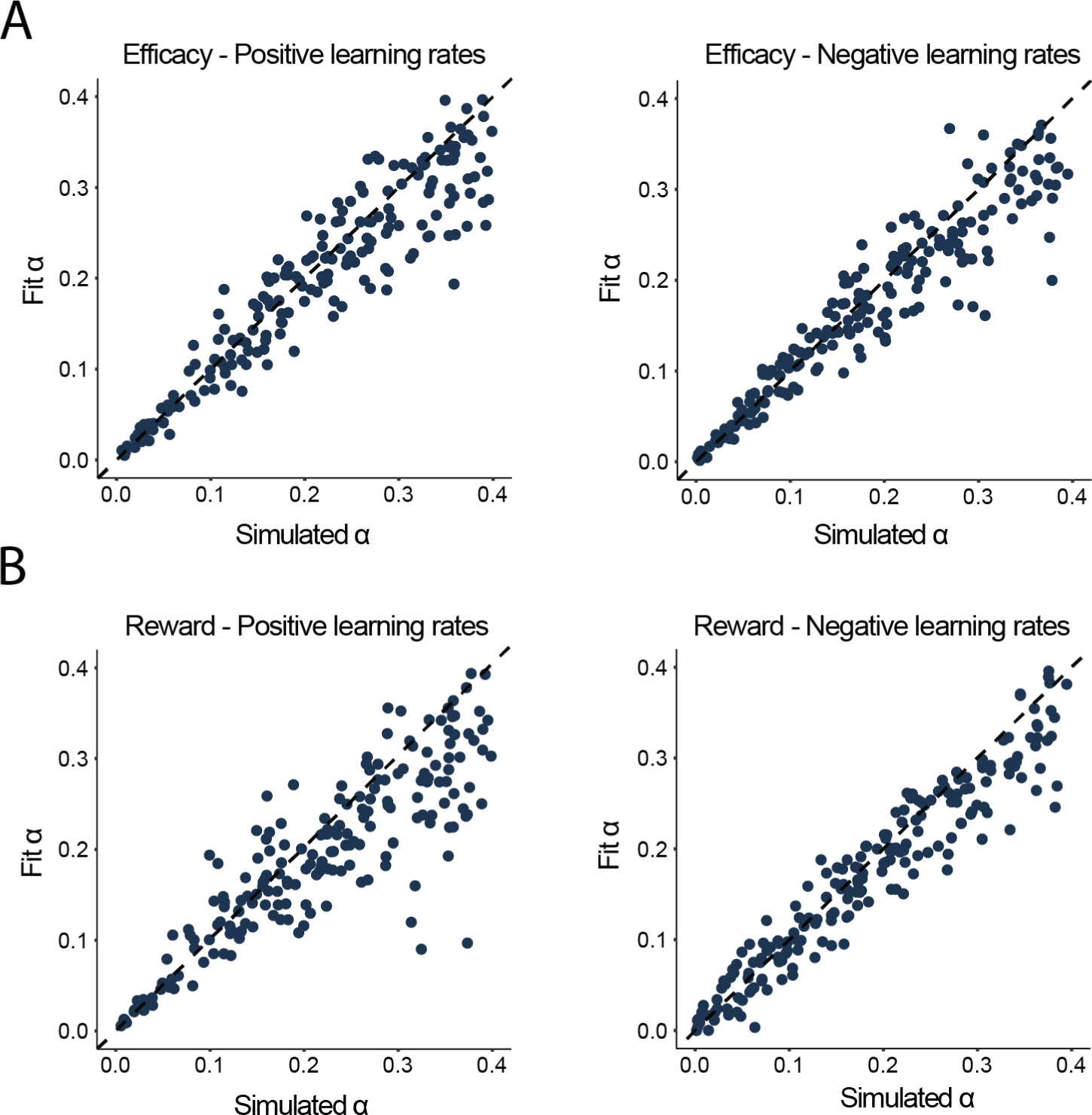
Parameter recovery study for the two learning rate models for the efficacy and reward learning. We simulated 200 subjects with the range of learning rates matched to the empirically observed range. The noise and the intercept parameters were fixed to match the empirically observed mean value. The number of trials and subjective estimates, as well as the sequence of efficacy and reward feedbacks were based on Experiment 1. **A. Learning rates for efficacy.** Simulated and recovered learning rates were highly correlated both in the case of positive (*r* = 0.93, *p*<0.001) and negative learning rates (*r* = 0.94, p<0.001). **B. Learning rates for reward.** Simulated and recovered learning rates were highly correlated both in the case of positive (*r* = 0.88, *p*<0.001) and negative learning rates (*r* = 0.96, *p*<0.001).

**Figure S2.**
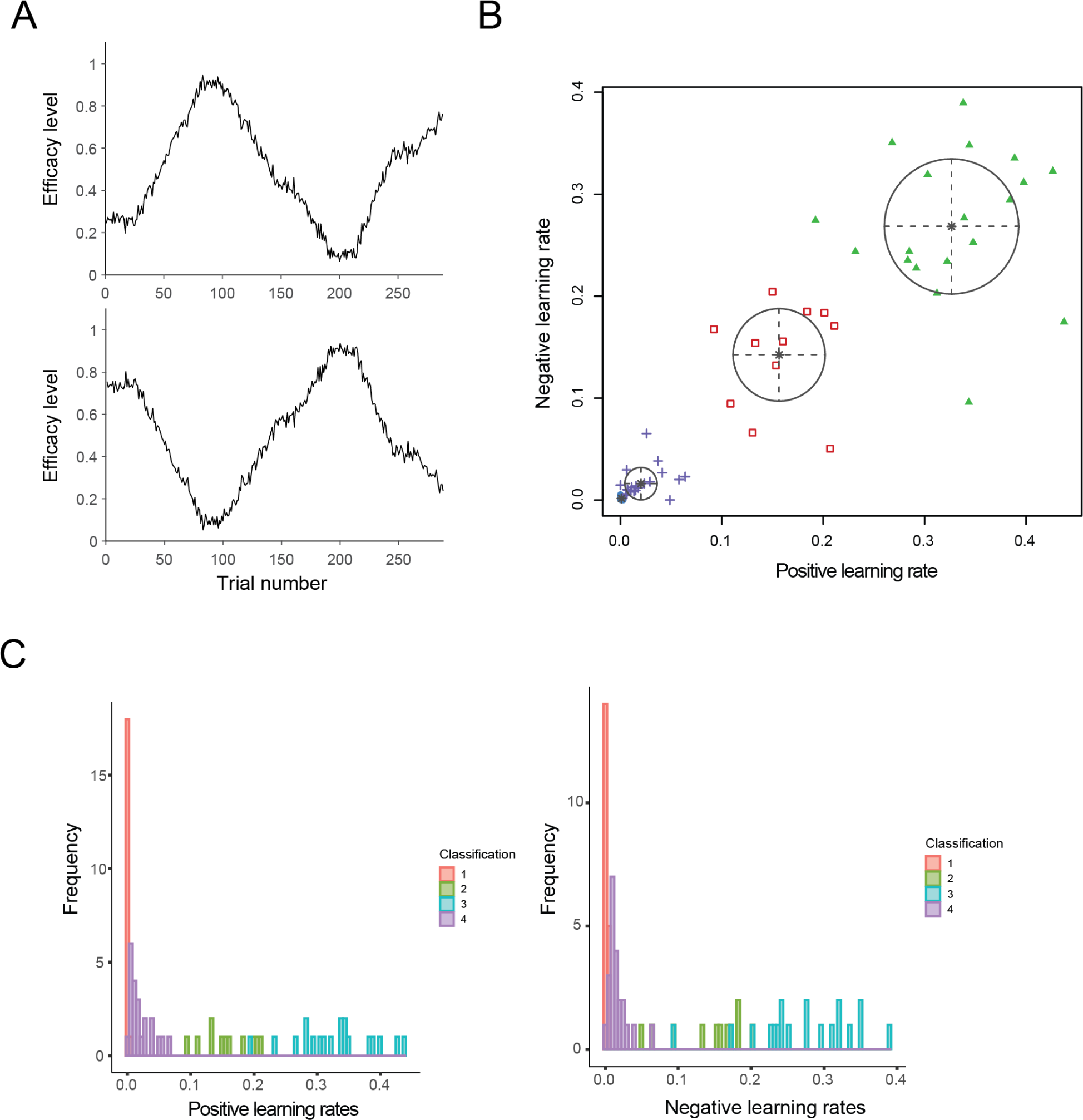
Efficacy drifts and the unsupervised clustering of the efficacy learning rates in Study 2. A. Efficacy drifts used in Study 2. The probability drifts of contingent feedback presented to the first (top) and second (bottom) half of the participants. B. The results of the winning Gaussian mixture model for clustering of the efficacy learning rates. One of the clusters (blue dots) included only the subjects with extremely small learning rates (all learning rates < 0.03; N=20; group 1 – blue dots). These subjects were excluded from the further analyses with the assumption that they did not pay attention to the feedbacks or that they were giving random answers to the probes. C. Histograms of the positive and negative learning rates and their clusters. For both then negative (left) and positive (right) learning rates cluster 1 included only the participants with very low learning rates.

**Figure S3.**
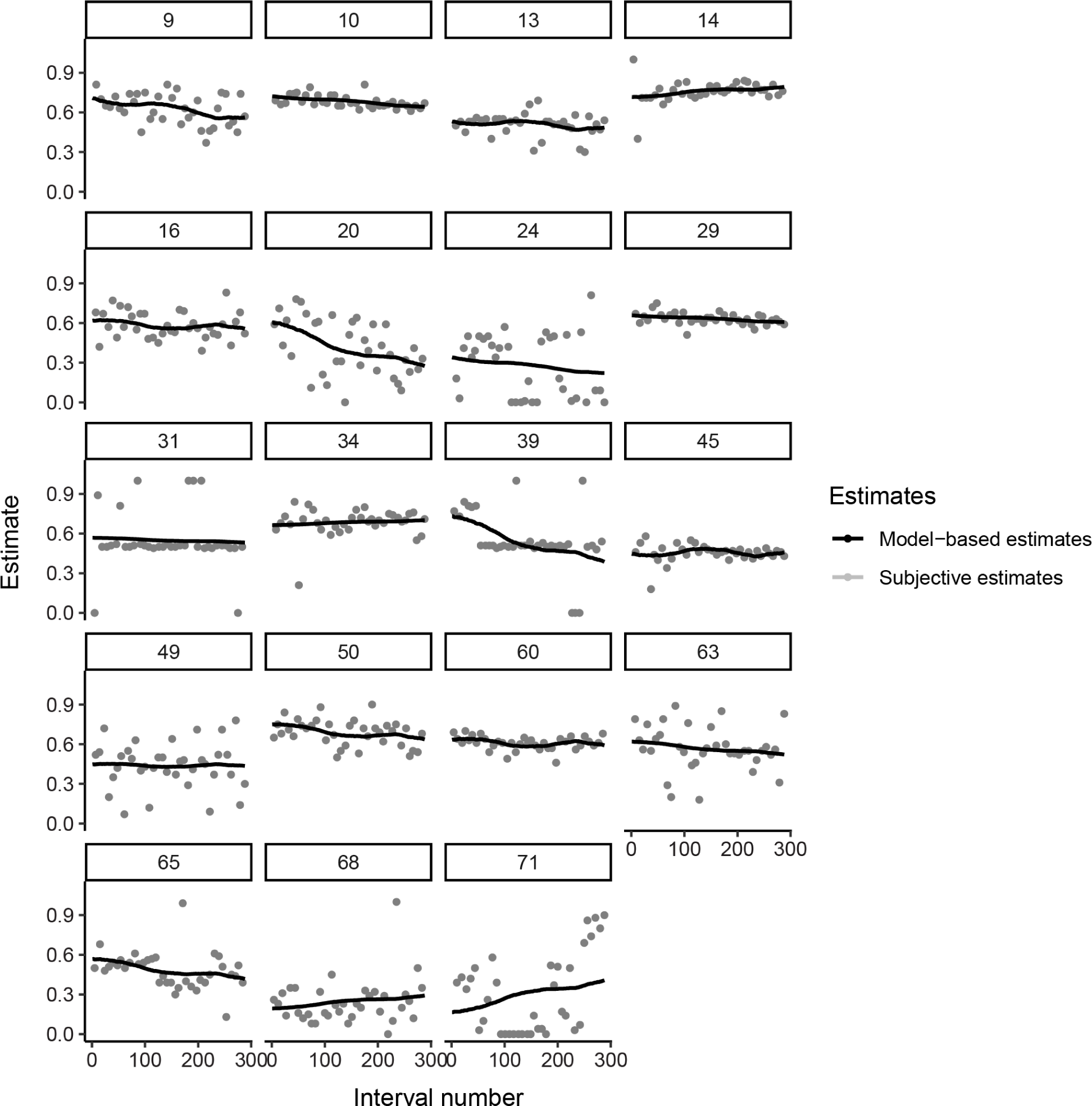
Self-reported and model-based efficacy estimates for subjects excluded from Study 2 based on very low learning rates. Subjective efficacy estimates (grey) and model-based efficacy estimates for each of the 19 subjects identified to form a cluster due to very low learning rates (both learning rates < 0.03) based on the unsupervised clustering algorithm using Gaussian mixture models. These subjects had low variance in their self-reported efficacy estimates, or did not appear to update their efficacy estimates based on feedbacks, suggesting that they were not paying attention to the feedbacks, or always providing very similar efficacy estimates.

**Figure S4.**
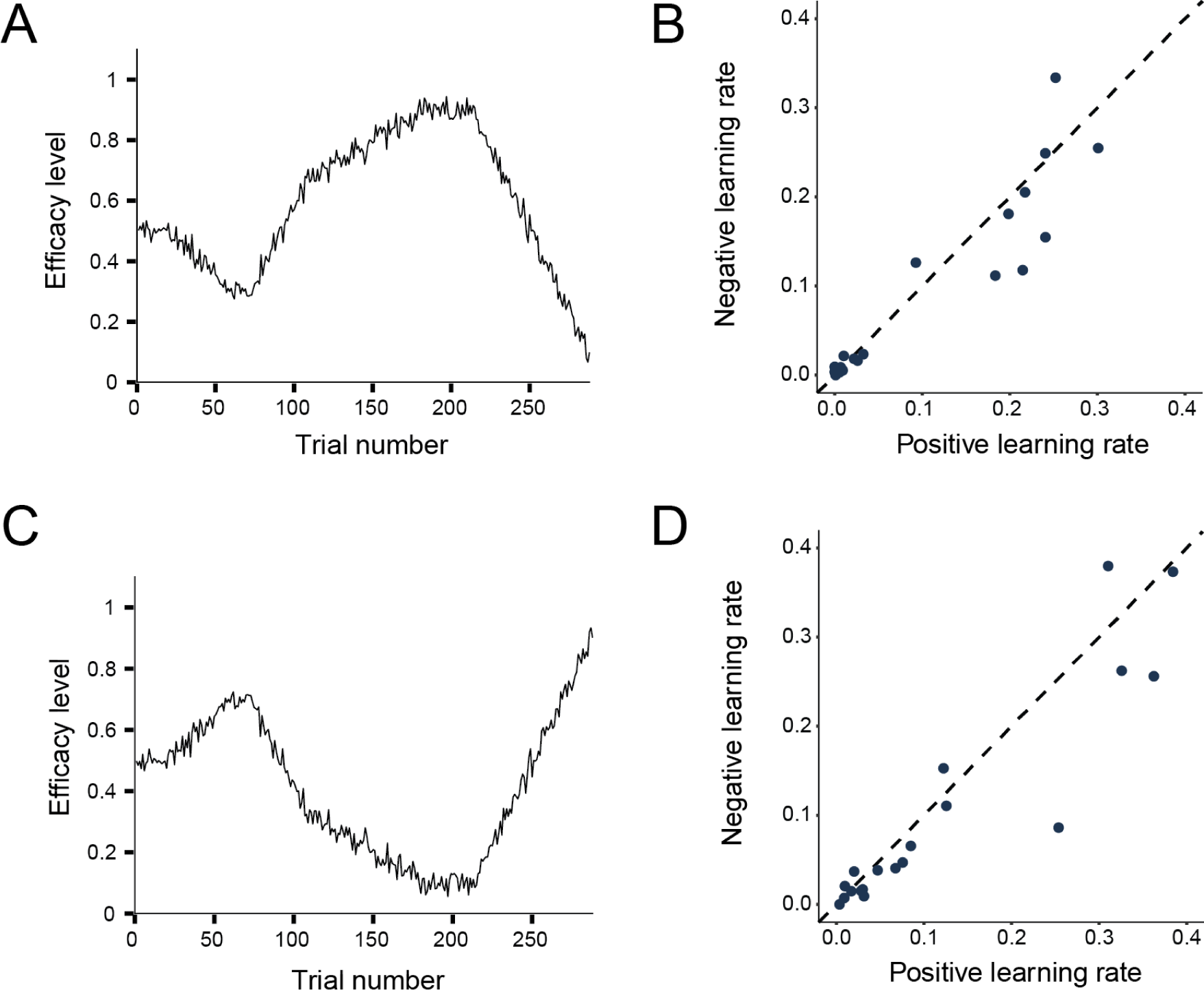
Efficacy learning in Study 1 – the effect of different feedback probability drifts on learning. **A.** The probability drift of contingent feedback presented to the first half of subjects. **B.** Learning rates for contingent and random feedback for the first half of subjects. Learning rates for the contingent and random feedback did not differ (*b =* 0.01; 95% CrI [-0.06, 0.08]; p*_b_* _< 0_ = 0.38; BF_10_ = 0.09). **C.** The inversed probability drift of contingent feedback presented to the second half of participants. **D.** Learning rates for contingent and random feedback did not differ (*b =* 0.02; 95% CrI [-0.06, 0.11]; p*_b_* _< 0_ = 0.31; BF_10_ = 0.10).

**Figure S5.**
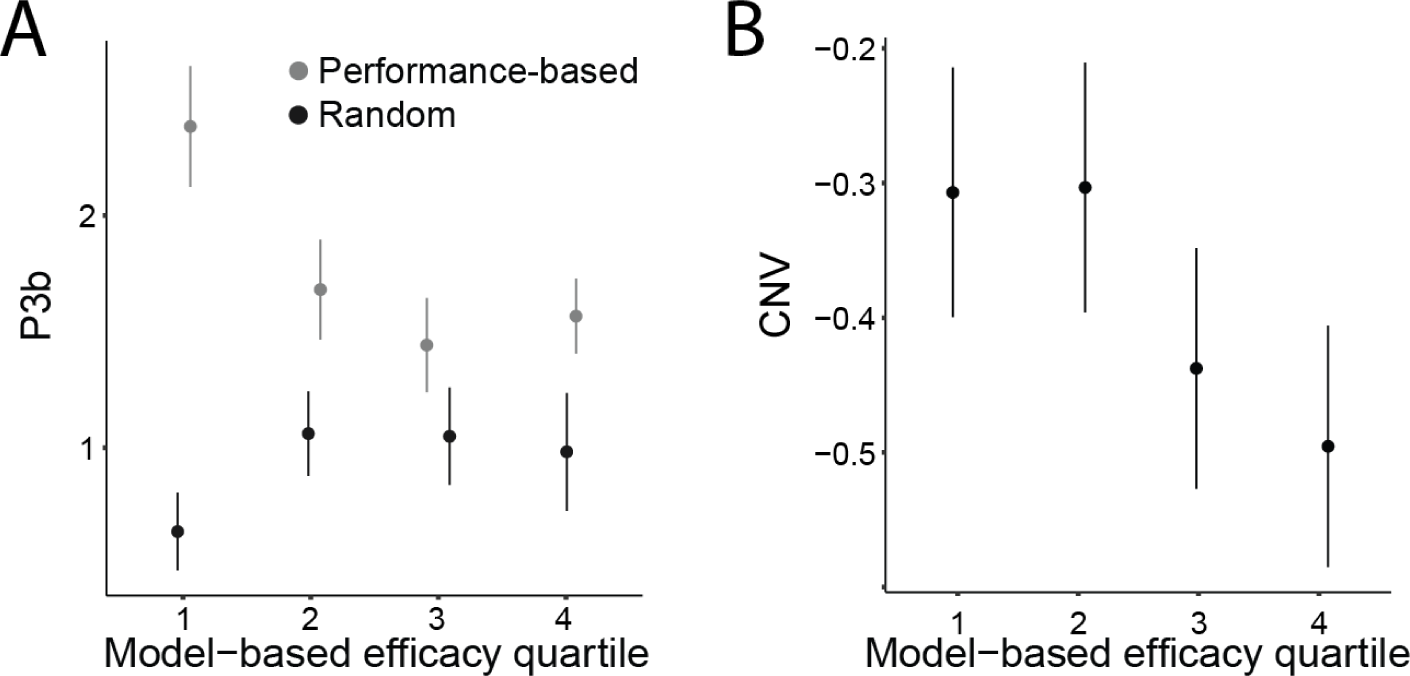
The influence of efficacy estimates on efficacy feedback processing and proactive cognitive control allocation. **A.** Means for the P3b amplitude in response to performance-based and random feedbacks across each quartile of the model-based efficacy estimates calculated for each subject. The P3b amplitude in response to performance-based feedback is higher when efficacy estimates are lower, suggesting more updating based on feedback. Error bars represent standard errors of the mean. **B.** Means for the CNV amplitude pre target onset across quartiles of the model-based efficacy estimates calculated for each subject. The CNV amplitude is more negative (more proactive control allocation) when efficacy estimates are higher. Error bars represent standard errors of the mean.

**Figure S6.**
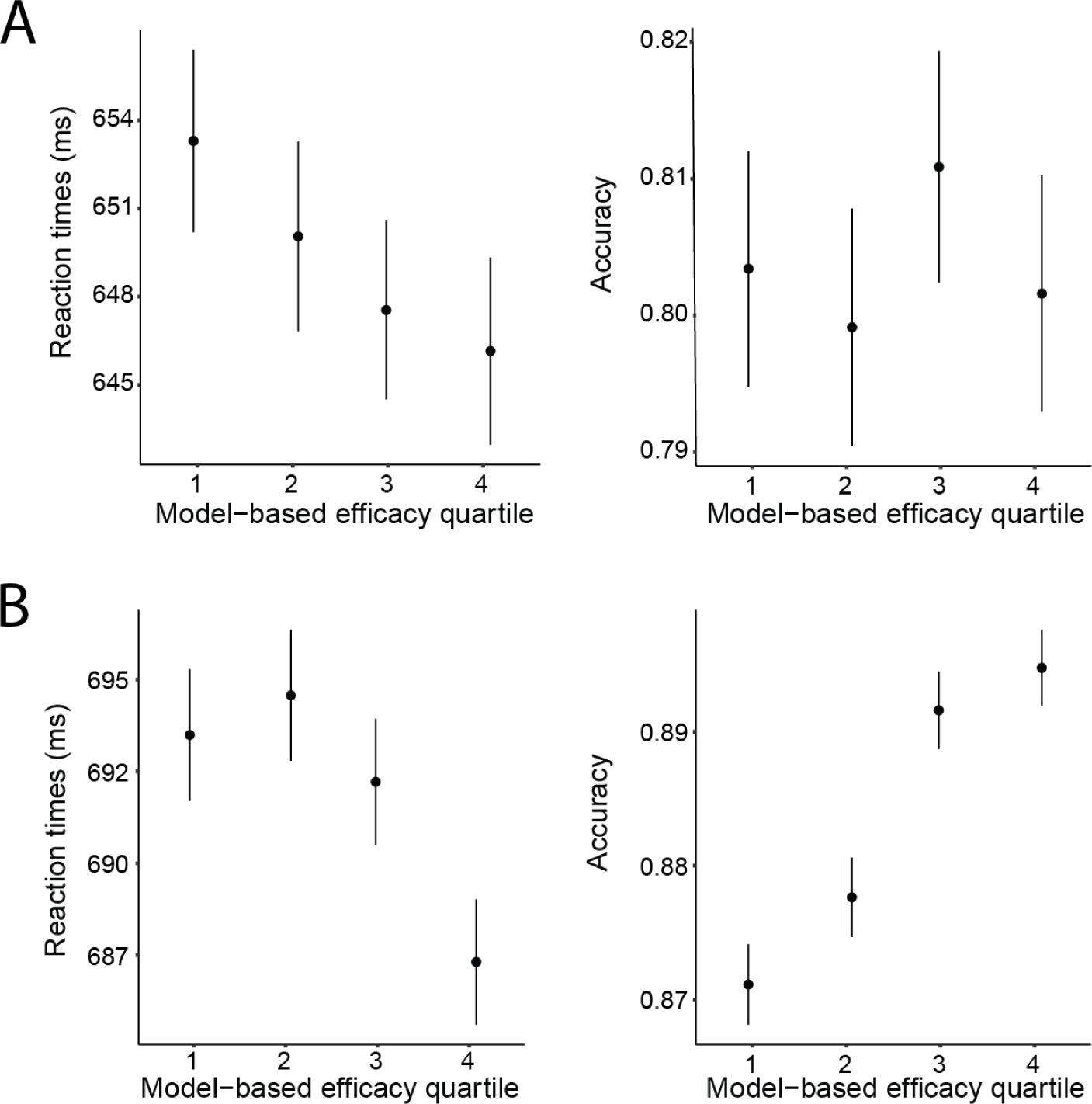
The influence of efficacy estimates on behavioral performance. **A.** Means for the reaction times (left) and accuracy (right) across quartiles of model-based efficacy estimates in Experiment 1. Reaction times are faster when efficacy estimates are higher, while there is no consistent pattern in accuracy data. Error bars represent standard errors of the mean. **B.** Means for the reaction times (left) and accuracy (right) across quartiles of model-based efficacy estimates in Experiment 2. Participants are faster to respond and more accurate when efficacy estimates are high relative to low. Error bars represent standard errors of the mean.

**Figure S7.**
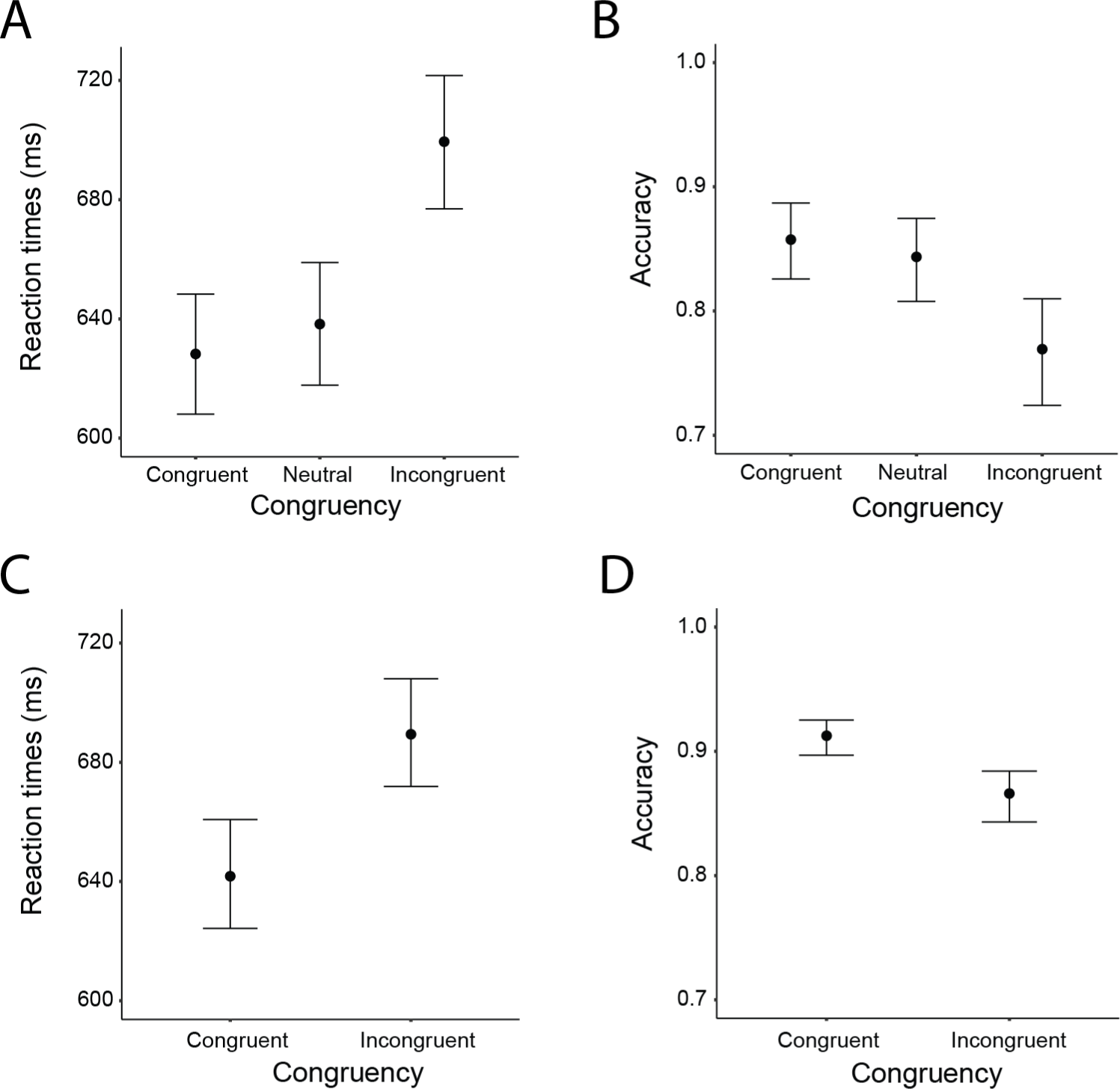
Behavioral effects of congruency in Study 1 and Study 2. **A.** In comparison to the neutral trials in Study 1, participants were faster to respond to congruent (*b* = 9.95; 95% CrI [3.33, 16.59]; p*_b_*_<0_ = 0.00; BF = 1.60) and slower to respond to incongruent trials (*b =* 61.24; 95% CrI [49.50, 72.59]; p*_b_* _< 0_ = 0.00; BF = 6.73). **B.** In comparison to the neutral trials in Study 1, participants were equally likely to be correct when responding to congruent (*b =* -0.11; 95% CrI [-0.29, 0.06]; p*_b_* _> 0_ = 0.091; BF = 0.01) and less likely to be correct when responding to incongruent trials (*b =* -0.48; 95% CrI [-0.64, -0.33]; p*b* > 0 = 0.00; BF > 8.22). The regression weights for the accuracy analysis are in log odds. **C.** In Study 2 participants were slower to respond to incongruent relative to congruent trials (*b =* 47.48; 95% CrI [40.69, 52.22]; p*_b_* _< 0_ = 0.001; BF = 50). **D.** In Study 2 participants were less accurate when responding to incongruent compared to congruent trials (*b =* -0.48; 95% CrI [-0.57, -0.39]; p*_b_* _< 0_ = 1; BF > 100).

**Figure S8.**
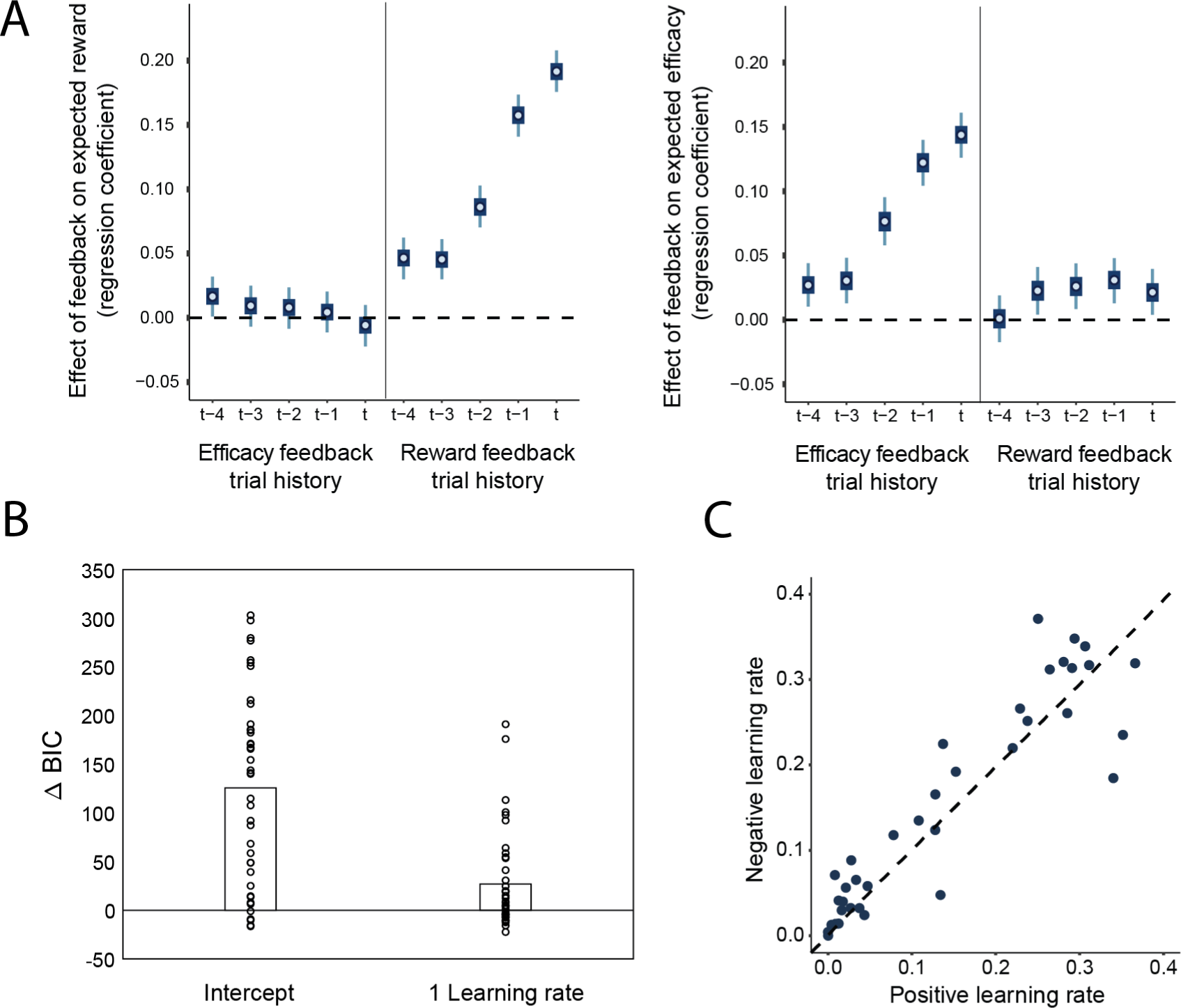
Reward rate learning – model-based analyses for Study 1. **A.** Regression weights for the influence of the current (t) and previous feedbacks on the subjective estimates of reward rate (left) and efficacy (right). Reward rate estimates are strongly predicted by the previous reward feedbacks, but not efficacy feedbacks. The reverse is true for the subjective efficacy estimates. Error bars represent 50% and 95% highest density intervals. **B.** Model comparison between the fitted learning models for the reward rate learning model. **C.** Positive and negative learning rate estimates for all subjects for the reward rate learning model. Negative learning rates were numerically, but not statistically larger than positive ones (*b =* 0.01; 95% CrI [-0.07, 0.09]; p*_b_* _< 0_ = 0.38; BF_10_ = 0.09)

## ERP analyses of the reward rate feedback in Study 1

To investigate the effect of the model-based reward rate estimates on the processing of reward, we performed complementary analyses on the P3b related to the processing of the reward feedback (Figure S9A). We expected that a similar learning mechanism should operate on both reward and efficacy feedback, but that the neural markers of feedback processing should be sensitive to different model-based estimates. Thus, we expected the reward-locked P3b to be sensitive to reward-rate estimates, but not to efficacy estimates.

Our results showed larger P3b amplitudes to no reward compared to reward feedback (note that reward rate was approximately .80 and negative feedback less expected overall), and the processing of the reward feedback was influenced by reward-rate estimates, although to lesser extent than for the P3b (Figure S9B; Table S6). Probing the effects of learning directly, we found robust effects of the prediction errors and learning rates (Figure S9C). The P3b to reward feedback was not influenced by the model-based efficacy estimates in either of these analyses.

---

**Figure S9.**
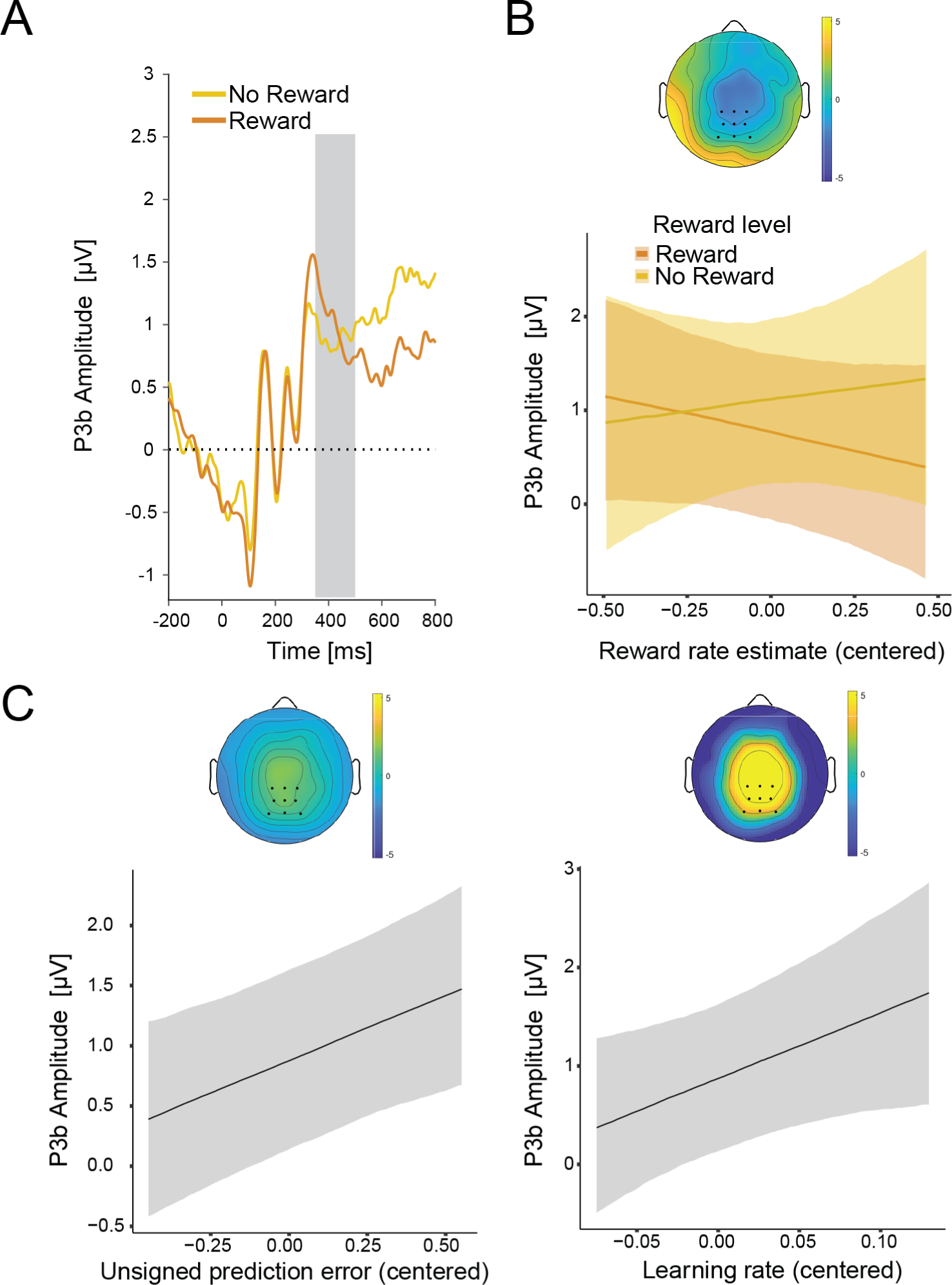
Learning reward rate estimate from reward feedback in Study 1. **A.** ERP average for the P3b locked to the onset of reward feedback. **B.** Interaction between reward feedback and the model-based reward rate estimate. **C.** The effects of unsigned prediction errors (left) and learning rates (right) on the P3b.

**Figure S10.**
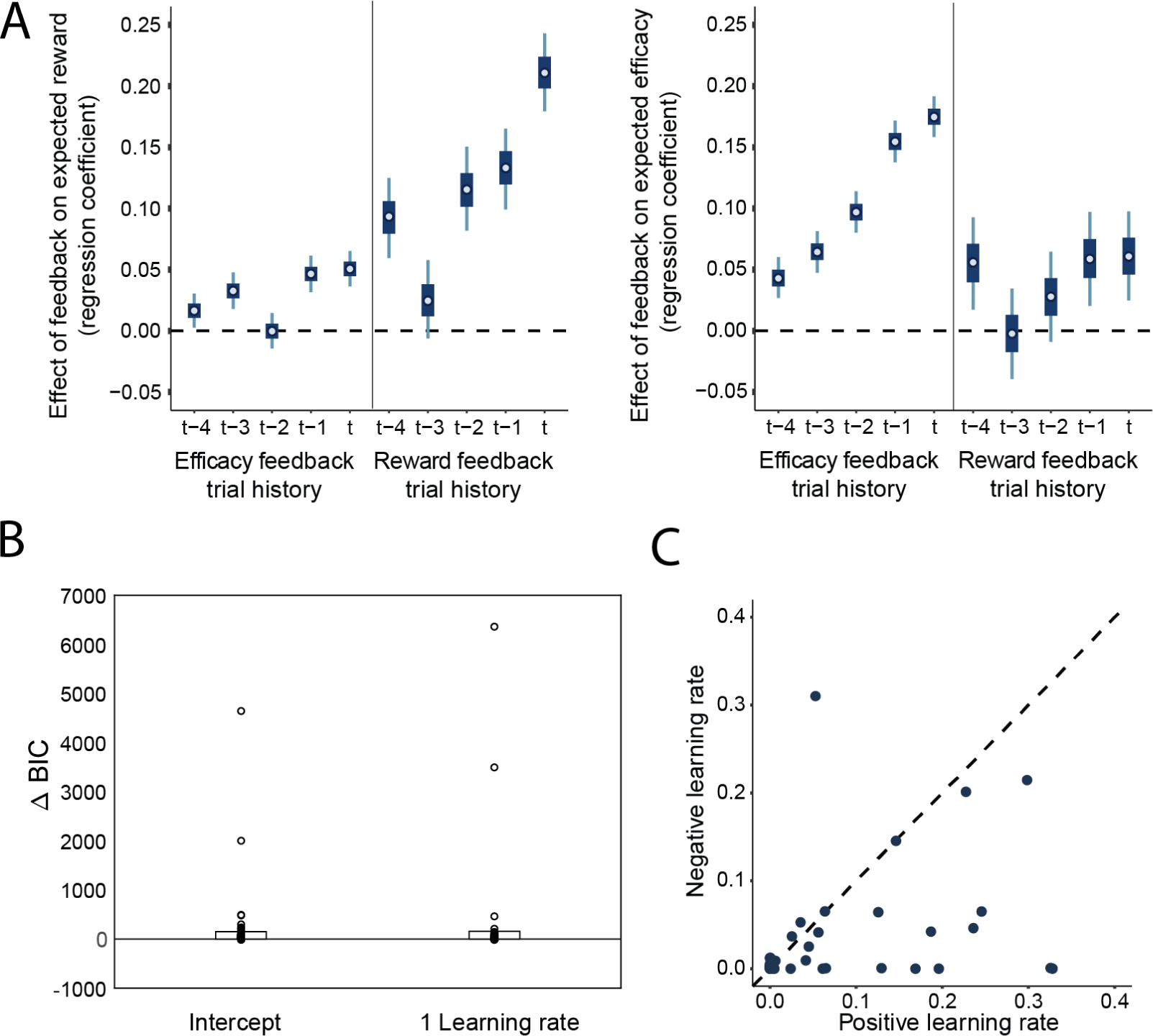
Efficacy and reward learning – model-based analyses for Study 2. **A.** Regression weights for the influence of the current (t) and previous feedbacks on the subjective estimates of reward rate (left) and efficacy (right). Reward rate estimates are strongly influenced by the previous reward feedbacks, and the reverse is true for the subjective efficacy estimates. Error bars represent 50% and 95% highest density intervals. **B.** Model comparison between the fitted learning models for the reward learning model. **C.** Positive and negative learning rate estimates for all subjects for the reward learning model. Positive learning rates were numerically, but not statistically larger than negative ones (*b =* 0.02; 95% CrI [-0.01, 0.05]; p*_b_* _< 0_ = 0.11; BF_10_ = 0.06)

**Table S1:** Study 1 regression weights for the models predicting subjective efficacy and reward rate estimates based on efficacy and reward feedback 5 trials back.

| *Parameter* | *Estimate* | *95% Credible interval* | *Posterior probability ($p < 0$)* | *BF<sub>10</sub>* |
| --- | --- | --- | --- | --- |
| *Subjective efficacy* |  |  |  |  |
| Intercept | 0.52 | 0.49, 0.55 | 0.00 |  |
| T1 Performance-Based - Random | 0.14 | 0.12, 0.16 | 0.00 | >100 |
| T2 Performance-Based - Random | 0.12 | 0.10, 0.14 | 0.00 | >100 |
| T3 Performance-Based - Random | 0.08 | 0.05, 0.10 | 0.00 | >100 |
| T4 Performance-Based - Random | 0.03 | 0.01, 0.05 | 0.00 | 2.20 |
| T5 Performance-Based - Random | 0.03 | 0.01, 0.05 | 0.00 | 1.72 |
| T1 Reward - No Reward | 0.02 | 0.00, 0.04 | 0.02 | 0.37 |
| T2 Reward - No Reward | 0.03 | 0.01, 0.05 | 0.00 | 4.23 |
| T3 Reward - No Reward | 0.03 | 0.01, 0.05 | 0.01 | 0.95 |
| T4 Reward - No Reward | 0.02 | 0.00, 0.04 | 0.02 | 0.39 |
| T5 Reward - No Reward | 0.00 | -0.02, 0.02 | 0.46 | 0.05 |
| *Subjective reward rate* |  |  |  |  |
| Intercept | 0.50 | 0.46, 0.54 | 0.00 |  |
| T1 Reward - No Reward | 0.19 | 0.17, 0.21 | 0.00 | >100 |
| T2 Reward - No Reward | 0.16 | 0.14, 0.18 | 0.00 | >100 |
| T3 Reward - No Reward | 0.09 | 0.07, 0.11 | 0.00 | >100 |
| T4 Reward - No Reward | 0.05 | 0.03, 0.06 | 0.00 | 30.86 |
| T5 Reward - No Reward | 0.05 | 0.03, 0.07 | 0.00 | >100 |
| T1 Performance-Based - Random | -0.01 | -0.03, 0.01 | 0.73 | 0.06 |
| T2 Performance-Based - Random | 0.00 | -0.01, 0.02 | 0.33 | 0.05 |
| T3 Performance-Based - Random | 0.01 | -0.01, 0.03 | 0.00 | 0.05 |
| T4 Performance-Based - Random | 0.02 | -0.01, 0.03 | 0.20 | 0.08 |
| T5 Performance-Based - Random | 0.02 | 0.00, 0.03 | 0.04 | 0.20 |

**Table S2:** Study 1 regression weights for the models predicting the P3b to efficacy feedback.

| *Parameter* | *Estimate* | *95%*<br>*Credible*<br>*interval* | *Posterior*<br>*probability*<br>*($p < 0$)* | *BF<sub>10</sub>* |
| --- | --- | --- | --- | --- |
| *Efficacy feedback* |  |  |  |  |
| Intercept | 1.38 | 0.67, 2.09 | 0.00 |  |
| Model-based efficacy | -0.29 | -1.56, 0.97 | 0.68 | 0.23 |
| Efficacy feedback type (PB-R) | 0.86 | 0.42, 1.31 | 0.00 | 30.86 |
| Model-based reward rate | -0.19 | -1.50, 1.18 | 0.62 | 0.23 |
| Model-based efficacy × Efficacy feedback type | -2.40 | -4.07, -0.74 | 1.00 | 24.40 |
| Model-based efficacy slope for Performance-Based feedback | -1.49 | -2.72, -0.27 | 0.97 | 1.52 |
| Model-based efficacy slope for Random feedback | 0.91 | -0.34, 2.21 | 0.12 | 0.44 |
| *Predictions errors and learning rates* |  |  |  |  |
| Intercept | 1.31 | 0.51, 2.09 | 0.00 |  |
| Unsigned prediction error | 1.25 | 0.35, 2.15 | 0.01 | 5.84 |
| Learning rate | 2.00 | -2.21, 6.04 | 0.17 | 1.08 |
| Model-based reward rate | -0.09 | -1.48, 1.34 | 0.55 | 0.23 |
| Unsigned prediction error × Learning rate | 0.57 | -4.87, 6.14 | 0.41 | 0.91 |

**Table S3:** Study 1 regression weights for the models predicting reaction times and accuracy based on efficacy and reward rate estimates.

| *Parameter* | *Estimate* | *95% Credible interval* | *Posterior probability (p<0)* | *BF<sub>01</sub>* |
| --- | --- | --- | --- | --- |
| *Reaction times* |  |  |  |  |
| Intercept | 655.37 | 635.09, 675.51 | 0.00 |  |
| Facilitation | 9.95 | 3.33, 16.59 | 0.00 | 1.60 |
| Interference | 61.24 | 49.50, 72.59 | 0.00 | 6.73 |
| Model-based efficacy | -16.00 | -34.91, 2.96 | 0.95 | 1.68 |
| Model-based reward rate | -1.56 | -27.66, 24.04 | 0.54 | 1.36 |
| *Accuracy* |  |  |  |  |
| Intercept | 1.56 | 1.33, 1.80 | 0.00 |  |
| Facilitation | -0.11 | -0.29, 0.06 | 0.91 | 0.01 |
| Interference | -0.48 | -0.64, -0.33 | 1.00 | 8.22 |
| Model-based efficacy | 0.12 | -0.20, 0.44 | 0.23 | 1.99 |
| Model-based reward rate | -0.36 | -0.73, 0.03 | 0.97 | 0.29 |

**Table S4:** Study 1 regression weights for the models predicting the CNV based on efficacy and reward rate.

| *Parameter* | *Estimate* | *95% Credible interval* | *Posterior probability (p<0)* | *BF<sub>01</sub>* |
| --- | --- | --- | --- | --- |
| *CNV* |  |  |  |  |
| Intercept | -0.37 | -0.58, -0.16 | 0.00 |  |
| Model-based efficacy | -0.35 | -0.85, 0.16 | 0.09 | 2.73 |
| Model-based reward rate | -0.09 | -0.72, 0.55 | 0.38 | 2.19 |

**Table S5:** Study 1 regression weights for the models predicting reaction times and accuracy based on the CNV.

| *Parameter* | *Estimate* | *95% Credible interval* | *Posterior probability ($p < 0$)* | *BF<sub>10</sub>* |
| --- | --- | --- | --- | --- |
| *Reaction times* |  |  |  |  |
| Intercept | 655.28 | 635.44, 675.91 | 0.00 |  |
| CNV amplitude | 11.41 | 8.09, 14.72 | 0.00 | >100 |
| *Accuracy* |  |  |  |  |
| Intercept | 1.33 | 1.11, 1.53 | 0.00 |  |
| CNV amplitude | -0.07 | -0.12, -0.01 | 0.99 | 3.14 |

**Table S6:** Study 1 regression weights for the models predicting the P3b to reward feedback.

| *Parameter* | *Estimate* | *95% Credible interval* | *Posterior probability (p<0)* | *BF<sub>10</sub>* |
| --- | --- | --- | --- | --- |
| *Efficacy feedback* |  |  |  |  |
| Intercept | 0.93 | 0.08, 1.74 | 0.02 |  |
| Model-based efficacy | 0.45 | -0.45, 1.32 | 0.16 | 0.15 |
| Reward feedback type (R-NR) | -0.35 | -0.90, 0.18 | 0.90 | 0.12 |
| Model-based reward rate | -0.15 | -1.55, 1.23 | 0.59 | 0.14 |
| Model-based reward × Reward feedback type | -1.26 | -3.73, 1.19 | 0.85 | 0.41 |
| Model-based efficacy slope for No Reward feedback | 0.48 | -1.34, 2.34 | 0.87 | 0.23 |
| Model-based efficacy slope for Reward feedback | -0.78 | -1.96, 0.39 | 0.32 | 0.21 |
| *Predictions errors and learning rates* |  |  |  |  |
| Intercept | 0.89 | 0.15, 1.64 | 0.01 |  |
| Unsigned prediction error | 1.09 | 0.38, 1.81 | 0.00 | 4.82 |
| Learning rate | 6.68 | -0.07, 12.99 | 0.03 | 4.72 |
| Model-based efficacy | 0.39 | -0.47, 1.24 | 0.19 | 0.13 |
| Unsigned prediction error × Learning rate | 0.94 | -8.06, 9.71 | 0.42 | 0.92 |

**Table S7:** Study 2 regression weights for the models predicting subjective efficacy and reward rate estimates based on efficacy and reward feedback 5 trials back.

| *Parameter* | *Estimate* | *95% Credible interval* | *Posterior probability ($p < 0$)* | *BF<sub>10</sub>* |
| --- | --- | --- | --- | --- |
| *Subjective efficacy* |  |  |  |  |
| Intercept | 0.43 | 0.37, 0.49 | 0.00 |  |
| T1 Performance-Based - Random | 0.17 | 0.16, 0.19 | 0.00 | >100 |
| T2 Performance-Based - Random | 0.15 | 0.13, 0.18 | 0.00 | >100 |
| T3 Performance-Based - Random | 0.10 | 0.08, 0.12 | 0.00 | >100 |
| T4 Performance-Based - Random | 0.06 | 0.04, 0.08 | 0.00 | >100 |
| T5 Performance-Based - Random | 0.04 | 0.02, 0.06 | 0.00 | >100 |
| T1 Reward - No Reward | 0.06 | 0.02, 0.10 | 0.00 | 5.42 |
| T2 Reward - No Reward | 0.06 | 0.01, 0.11 | 0.01 | 2.23 |
| T3 Reward - No Reward | 0.03 | -0.02, 0.07 | 0.11 | 0.23 |
| T4 Reward - No Reward | 0.00 | -0.05, 0.04 | 0.54 | 0.11 |
| T5 Reward - No Reward | 0.06 | 0.01, 0.10 | 0.01 | 1.69 |
| *Subjective reward rate* |  |  |  |  |
| Intercept | 0.36 | 0.29, 0.43 | 0.00 |  |
| T1 Reward - No Reward | 0.21 | 0.17, 0.25 | 0.00 | >100 |
| T2 Reward - No Reward | 0.13 | 0.09, 0.17 | 0.00 | >100 |
| T3 Reward - No Reward | 0.12 | 0.07, 0.16 | 0.00 | >100 |
| T4 Reward - No Reward | 0.03 | -0.01, 0.06 | 0.09 | 0.21 |
| T5 Reward - No Reward | 0.09 | 0.05, 0.13 | 0.00 | >100 |
| T1 Performance-Based - Random | 0.05 | 0.03, 0.07 | 0.00 | 0.06 |
| T2 Performance-Based - Random | 0.05 | 0.03, 0.06 | 0.00 | 0.05 |
| T3 Performance-Based - Random | 0.00 | -0.02, 0.02 | 0.51 | 0.05 |
| T4 Performance-Based - Random | 0.03 | 0.02, 0.05 | 0.00 | 0.08 |
| T5 Performance-Based - Random | 0.02 | 0.00, 0.03 | 0.03 | 0.20 |

**Table S8:** Study 2 regression weights for the models predicting reaction times and accuracy based on efficacy and reward rate estimates. Bayes Factors calculated only for the parameters with informative priors. Posteriors from Study 1 were used as priors for the effects of congruency, efficacy, and reward.

| *Parameter* | *Estimate* | *95% Credible interval* | *Posterior probability ($p < 0$)* | *BF<sub>01</sub>* |
| --- | --- | --- | --- | --- |
| *Reaction times* |  |  |  |  |
| Intercept | 643.02 | 625.42, 661.81 | 0.00 |  |
| Incongruent-Congruent | 47.48 | 40.69, 55.22 | 0.00 | 0.02 |
| Model-based efficacy | -10.25 | -19.86, -0.20 | 0.98 | 2.33 |
| Model-based reward | -0.17 | -21.15, 20.34 | 0.51 | 1.24 |
| Interval length | -1.76 | -2.73, -0.83 | 1.00 |  |
| Interval congruency | -5.34 | -6.69, -4.00 | 1.00 |  |
| *Accuracy* |  |  |  |  |
| Intercept | 2.35 | 2.16, 2.52 | 0.00 |  |
| Incongruent-Congruent | -0.48 | -0.57, -0.39 | 1.00 | >100 |
| Model-based efficacy | 0.23 | 0.03, 0.43 | 0.01 | >100 |
| Model-based reward | -0.29 | -0.57, -0.01 | 0.98 | 1.07 |
| Interval length | -0.13 | -0.16, -0.10 | 1.00 |  |
| Interval congruency | 0.04 | 0.00, 0.08 | 0.03 |  |

**Table S9:** Drift Diffusion model estimates.

| *Parameter* | *Estimate* | *95% Credible interval* | *Posterior probability ($p < 0$)* |
| --- | --- | --- | --- |
| *Drift rate* |  |  |  |
| Congruent-Incongruent | 0.82 | 0.74, 0.92 | 0.00 |
| Model-based efficacy | 0.29 | 0.14, 0.40 | 0.00 |
| Model-based reward | 0.20 | -0.39, 0.04 | 0.50 |
| *Threshold* |  |  |  |
| Model-based efficacy | -0.00 | -0.04, 0.04 | 0.57 |
| Model-based reward rate | -0.04 | -0.14, 0.05 | 0.81 |

## Acknowledgements

This work was supported by the Special Research Fund (BOF) of Ghent University [grant #01D02415] (I.G.), the Research Foundation Flanders (FWO) travel grant [grant #V432718N] (I.G.), a Center of Biomedical Research Excellence grant P20GM103645 from the National Institute of General Medical Sciences (A.S.), the Alfred P. Sloan Foundation Research Fellowship in Neuroscience (A.S.), and an NSF Graduate Research Fellowship (M.P.F). The funding sources were not involved in the study design; collection, analysis, and interpretation of data; writing of the report; and decision to submit the article for publication. We would like to thank Natalie Knowles and Hattie Xu for help with data collection for Study 1, Peyton Strong for help with data collection for Study 2, Carolyn Dean Wolf and Liz Cory for help with programming the Study 1 task, Harrison Ritz for help with constructing the efficacy drifts, and Xiamin Leng for advice on fitting the drift-diffusion models.

## Data availability

Analysis scripts are available on this link: https://github.com/igrahek/LFXC_EEG_2022.git Please contact the authors for raw data.

## Footnotes

1 Note that the prior distributions are set in log-odds.

2 Note that the CNV is a negative component, thus higher CNV amplitudes (i.e., more control allocation) are more negative.

## Notes

Conflicts of interest: The authors have no conflicts of interest to declare.

### Competing Interest Statement

The authors have declared no competing interest.

### Summary of Updates

We included a parameter recovery study, clarifications of the goals of the study, and more details on model fitting.

